# Population-scale gene expression analysis reveals the contribution of expression diversity to the modern wheat improvement

**DOI:** 10.1101/2025.02.18.638840

**Authors:** Zhimeng Zhang, Shengwei Ma, Mou Yin, Caihong Zhao, Xinyu Zhao, Yang Yu, Haojie Wang, Xuanzhao Li, Yaoqi Si, Jianqing Niu, Jingzhong Xie, Limin Wang, Jiajie Wu, Yanming Zhang, Qi Zheng, Shusong Zheng, Ni Jiang, Xigang Liu, Hong-Qing Ling, Fei He

## Abstract

Changes in gene expression are crucial for crop breeding, yet population genomics has primarily focused on sequence polymorphisms rather than gene expression diversity. The strategy of using single genome reference for RNA-seq analysis could not handle introgression bias, especially for hexaploidy wheat. Here, we conducted RNA-seq for 328 wheat lines, including representative diverse landraces and elite cultivars from China and the United States, to investigate the role of gene expression variation in shaping agronomic traits. Using pan-genome resources, we identified 20,615 more transcripts than using the ‘Chinese Spring’ reference genome alone. We constructed a pan-gene atlas regulatory map through eQTL analysis, demonstrating that genes introgressed from wild relatives were under tight genetic control. Genes responding to environmental stress show higher activity after introgressed into the wheat genome, demonstrating how long-term breeding selection impacted the gene expression regulation of targeted introgression. Multi-omics modeling identified 231 high-confidence candidate genes for 34 field agronomic traits and the seedling resistance phenotypes of 8 powdery mildew isolates. More than one fifth of those candidates have no homolog in ‘Chinese Spring’ reference genome. By utilizing the indexed KN9204 EMS library, 80% candidates showed significant trait difference between wild type and mutant lines. Furthermore, directional shifts in genes of which expression were changed by breeding improvement demonstrated distinct adaptations to local environments. Our study constructed a pan-gene atlas to correct the reference bias of reads mapping in RNA-seq studies and revealed the expression patterns of introgressed genes within the wheat genome and their regulatory mechanisms, which highlighted the impact of breeding selection on gene expression of the world’s most important crop.

## Introduction

Wheat is the most important food crop worldwide, supplying 19% of the daily caloric intake and 21% of the protein needs for people [1]. Scientists have successfully developed high-yield, disease-resistant, and stress-tolerant wheat varieties in modern breeding. This genetic improvement is particularly essential when facing the dual challenges of climate change and a growing global population [2]. Wheat is vulnerable to various diseases, including *Fusarium* head blight, rust, and powdery mildew, in different growing environments. Genetic diversity studies enable breeders to identify and utilize disease resistance genes, leading to the development of wheat accessions with enhanced resistance. Moreover, genetic diversity offers a broad gene pool for wheat to adapt to harsh climate conditions, such as drought and saline soils [3]. Maintaining and expanding wheat genetic diversity is key to preventing genetic bottlenecks and provides abundant genetic resources for future breeding [4].

The importance of DNA diversity (i.e., single nucleotide polymorphism, SNP) were well studied in recent years. One of the conclusions is that the introgression of landraces and wild relatives has shaped the genetic diversity of wheat. As a rich genetic resource, landraces have been less influenced by historical and geographical effects, preserving a significant number of genes that have not been widely utilized in modern breeding. These genes can be used to improve the diversity of cultivars, particularly in complex quantitative traits and stress-resistance traits. *Cheng et al.* conducted whole-genome resequencing of 827 A. E. Watkins landraces and 208 modern varieties to investigate the genetic and phenotypic diversity present in the historical Watkins germplasm collection. The research highlighted the unique allelic and haplotype variations in landraces, thereby providing novel resources for future breeding efforts [5]. *Niu et al*. collected 180 landraces and 175 cultivars to investigate the genetic variation in modern Chinese and American breeding programs through whole-genome resequencing. The study highlighted the necessity of conserving and utilizing the genetic diversity of landraces during breeding [6]. Additionally, the introgression of wild relatives serves as a potential resource for enhancing genetic diversity. *He et al.* performed exome sequencing on roughly 1,000 hexaploid and tetraploid wheat lines, identifying gene introgression from wild relatives, and highlighting the important contribution of historic gene flow from wild relatives to the adaptive landscape of modern bread wheat [7]. A majority of wild relatives from the *Triticeae* tribe can hybridize with wheat, and through backcrossing or chromosome engineering, chromosome segments carrying specific alleles can be introduced into wheat. To date, numerous genes, particularly disease resistance genes, have been transferred into wheat from rye, various *Triticum* species, *Aegilops*, *Thinopyrum* and *Dasypyrum* genera [8–13]. However, only a small portion of the existing genetic diversity has been utilized, with many genes and alleles yet to be exploited for broader trait improvements [14].

Studies on how breeders’ selection impacted the gene expression landscape at a genome-wide level is lacking, although studies of individual genes showed the crucial role of gene expression regulation to wheat improvement. For example, the regulatory expression of *Vernalization 1* (*VRN1*), *Vernalization 2* (*VRN2*), and *Vernalization 3* (*VRN3*)/*Flowering locus T1* (*FT1*) during the vernalization process is essential for determining flowering time and environmental adaptability in wheat [15]. The photoperiod-insensitive *Ppd-1* allele (e.g, *Ppd-D1a*) is widely used in breeding to reduce the requirement for long daylight hours. The *Ppd-1* gene regulates the expression of the flowering activator *VRN3*/*FT1*, promoting earlier flowering under shorter photoperiods, which enhances grain formation and adaptive growth in wheat [16–19]. The upregulation of the NAM-ATAF-CUC transcription factor *TaNAC100* facilitates the expression of starch synthesis-related genes, *TaGBSS1* and *TaSUS2*, increasing starch content in seeds. Furthermore, the overexpression of *TaNAC100* also impacts the total protein content in wheat seeds, playing a crucial role in balancing seed storage proteins and starch [20]. Nitrate-induced NAC transcription factor *TaNAC2*-*5A* positively regulates the expression of *TaNRT2.5* and *TaNRT2.1*. Overexpression of *TaNAC2*-*5A* significantly enhances nitrate absorption, grain nitrogen concentration, and wheat yield, indicating its potential in simultaneously improving wheat yield and grain protein content [21]. Under conditions of salt stress, wheat lines characterized by high expression levels of the *TaSOS1* gene display more robust root systems and enhanced water potential, which correlates with improved salt tolerance. This suggests that *TaSOS1* holds considerable potential for application in salt tolerance breeding [22, 23]. Although functional studies of these genes have highlighted the significance of gene expression regulation in wheat improvement, a significant gap remains in understanding how breeding selection systematically affects gene expression at the genome-wide level. Further exploration in this field will provide new directions for optimizing wheat breeding strategies.

A special case of dysregulation in homoeologs was discovered to be correlated with agronomic traits of wheat, which was later proved to be due to reference bias [24, 25]. Reference bias refers to the underestimation of transcripts from non-reference alleles during quantification, which can impact the accuracy of subsequent conclusions. This bias is particularly pronounced in complex polyploid genomes, such as hexaploid wheat, which often contain highly heterologous gene blocks resulting from the introgression of wild relatives. Consequently, they proposed a method to reduce reference bias by constructing a pan-transcriptome reference that integrates the gene models from Chinese Spring (CS) and nine genomes from the 10+ wheat genomes project. They incorporated only the transcripts of genes with a 1-to-1 orthologous relationship to the genes of Chinese Spring, subsequently merging the quantitative results based on the homology relationships of Chinese Spring [25]. Their method primarily addressed errors caused by allelic variation, however, they still face limitations regarding presence/absence variations.

Despite the abundant genome assemblies of wheat cultivars and relative species, RNA-seq of cultivated wheat still uses gene models mostly from one single reference genome, i.e., Chinese Spring. Here we utilized 24 genome assemblies to build a non-redundant gene atlas along with the genes of Chinese Spring. We selected 328 hexaploid common wheat accessions from earlier resequencing studies [6], and performed RNA sequencing. We aligned RNA-seq reads with the pan-gene atlas and integrated high-density resequencing SNP data to construct a genetic regulatory map of gene expression. Introgressed genes were found to be *trans*-regulated, with resistance genes enriched in introgression lines. By analyzing 34 field agronomic traits and the seedling resistance phenotypes of 8 powdery mildew *Blumeria graminis* f. *sp*. *tritici* (*Bgt*) isolates, we identified 231 high-confidence candidate genes, including 54 non-CS genes, and validated 58 agronomic trait-associated genes using the KN9204 mutant library. Differentially expressed genes between cultivars and landraces revealed divergent breeding directions in different countries, with regulatory regions under stronger selection. Modern breeding significantly altered regulatory networks in cultivars compared to landraces. This study highlights the genetic regulation of the wheat transcriptome and its impact on breeding, providing valuable resources for wheat breeding improvement.

## Results

### The impact of single reference in RNA-seq reads mapping leads to underestimate of expression for introgression

To investigate the genetic regulation of gene expression in modern wheat breeding improvement, we performed RNA sequencing (RNA-seq) on 2-week-old seedlings from a panel of 328 wheat lines (Supplementary Table 1), including 172 representative diverse landraces (LR) and 92 modern Chinese cultivars (MCC) and 64 modern United States cultivars (USMC), which had been previously subjected to whole-genome resequencing [6]. An average of 77.0 million paired-end Illumina reads (2×150 bp) was obtained per sample, followed by quality control and alignment to the CS RefSeq v1.1 reference genome.

The presence of introgressions can lead to reference bias in wheat RNA-seq analysis [25]. So we first tested the effect of reference bias for the most widely deployed 1RS.1BL wild relative introgression, where the long arm of *Secale cereal* (rye) chr1R translocated with the long arm of wheat chr1B 50 lines of our panel carried this introgression (Supplementary Table 2). The number of expressed genes in the sliding window along the CS chromosome arm are lower for lines with introgression (Fig. 1a). On average, each 10 Mb chromosome window has 24 and 26 expressed genes for introgressed and non-introgressed lines, respectively (*p*-value < 2.22×10^-16^) (Fig. 1a). The average expression levels of 1RS.1BL translocation lines were lower than that of non-introgression 1BS arm (*p*-value < 2.22×10^-16^) (Fig. 1b and Supplementary Fig. 1). Since the coding sequence of rye has diverged with diploid wheat 9.6 million years ago [26], quantifying gene expression for the rye introgression based on the standard reference Chinese Spring may underestimate the gene expression levels.

**Fig. 1.**
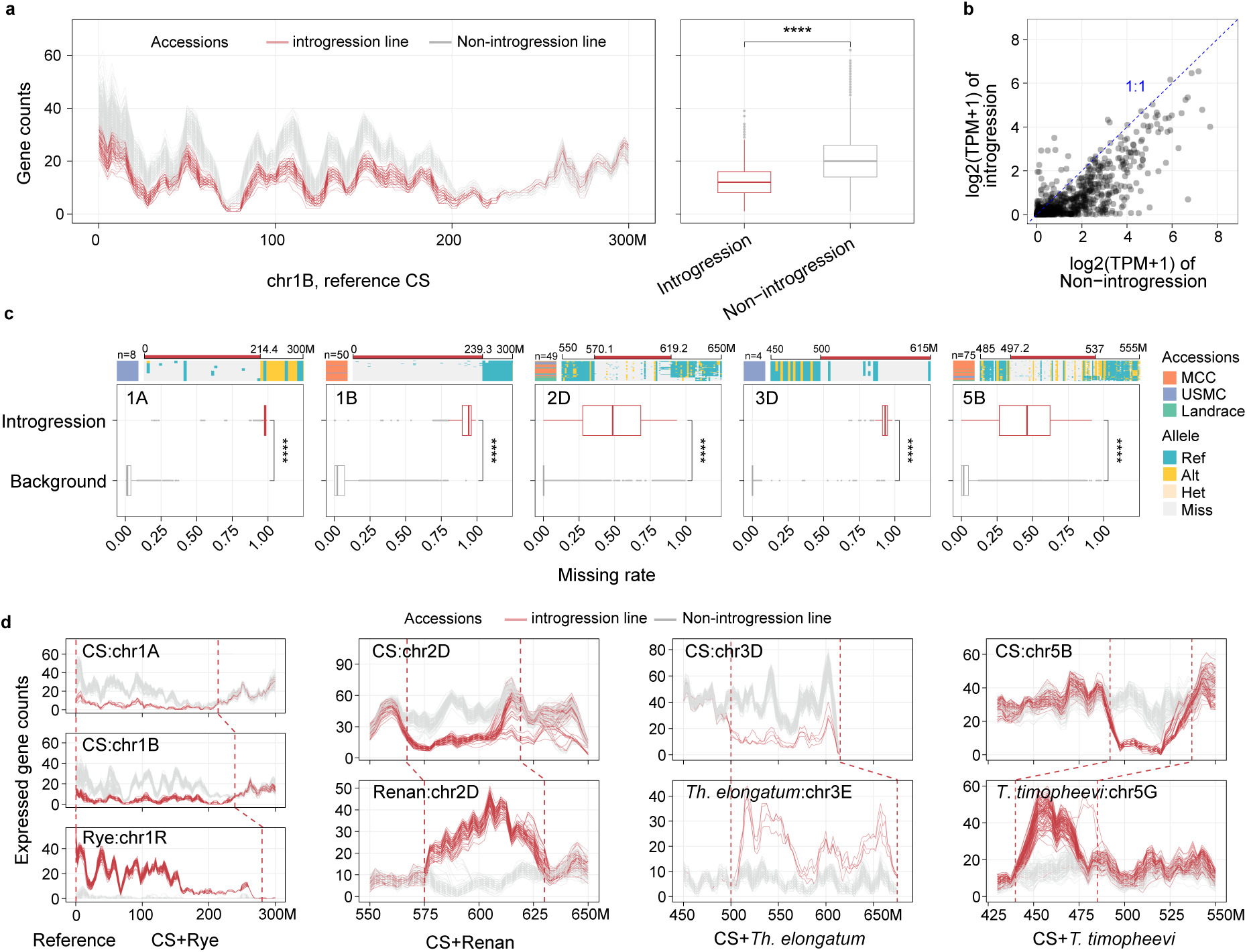
Assessing reference bias in alien introgression of wheat. **a** The left panel shows the number of expressed genes (TPM > 0.5) in introgression and non-introgression lines along chromosome 1BS (0-300 Mb), based on Chinese Spring RefSeq v1.1 as the reference genome, using a sliding window of 10 Mb and a step size of 2.5 Mb. The right panel presents boxplots comparing the number of expressed genes between introgression and non-introgression lines within the chromosome 1BS (0-280 Mb) (Wilcoxon rank sum test, \*\**p* < 0.01; \**p* < 0.05). **b** Average gene expression of shared genes between introgression and non-introgression lines in chromosome 1BS (0-280 Mb) region, with Chinese Spring RefSeq v1.1 as the reference genome. **c** A heatmap displays missing genotype sites and a boxplot illustrates the genotype missing rate with the Chinese Spring RefSeq v1.1 as the reference genome. In the heatmap, n represents the number of accessions, orange represents MCC, cornflower blue represents USMC, and green represents Landrace. The upper red color bar indicates introgressed fragments, and four genotype categories are depicted: blue-green for homozygous reference alleles, gold for homozygous alternative alleles, light gold for heterozygous alleles, and light gray for missing genotypes. The lower boxplot represents genotype missing rates calculated within a 2 Mb sliding window and a 1 Mb step size. **d** The number of expressed genes was calculated using a merged reference genome of Chinese Spring and introgression donors. The method for quantifying expressed genes is consistent with that in Fig. 1a, non-introgression lines include only the 100 samples with the most consistent gene expression (gray lines)The upper line chart shows the number of expressed genes from Chinese Spring in the merged genome, while the lower line chart displays the number of expressed genes from the introgressed reference genome in the merged assembly.

In order to understand the scale of reference bias due to large segments introgression, we first need to know where those large introgression locate. We utilized the genotype missing rate to identify multiple deletion fragments across the CS genome. Those regions with continuous high missing rate were considered as potential introgression [27]. We focused on five introgressions larger than 20 Mb from known donors, including 1RS.1AL (chr1A: 0-214.4 Mb) and 1RS.1BL (chr1B: 0-239.3 Mb) [28], as well as chr2D: 570.1-619.2 Mb from *Aegilops markgrafii* [29, 30], chr3D: 500-615.5 Mb, and the terminal ∼60 Mb of chr3D in *LongReach Lancer*, confirmed as a *Thinopyrum ponticum* introgression [29]. Additionally, chr5B: 497.2-537 Mb originates from *Triticum timopheevii* (Fig. 1c and Supplemental Table 2) [27]. Those regions are introgression rather than chromosome deletion due to 1) the missing rate is significantly higher than the other genomic regions (Fig. 1c); and 2) missing rate are lower in protein coding sequences than non-coding regions (Supplementary Fig. 3), indicating the presence of the homologs in the collinear regions between CS and nonCS. The expression of these introgressed fragments was also underestimated (Supplementary Fig. 1, 2).

To accurately quantify gene expression in introgression fragments, we merged the donor genome with the Chinese Spring reference genome (see Methods) and performed expression analysis for all wheat samples. Non-introgression lines exhibited a higher number of expressed genes on the Chinese Spring chromosomes compared to introgression lines. Conversely, in introgression lines, actively expressed genes were recovered in the donor genome within each 10 Mb window (Fig. 1d and supplemental Note). Thus, the reference bias can be severe when measuring expression using RNA-seq data for wheat lines carrying introgression. Gene model of donor species are needed for accurately estimate the expression of introgressions other than using a single reference genome.

### Utilizing pan-genome resource for accurate quantification of gene expression

Recent studies showed wild relative such as wild emmer introgression can contribute up to 15% of the wheat genome and most introgressed fragments are less than 1 Mb in size [7, 31]. In addition to the large introgressed fragments from Rye, *Ae. markgrafii*, *T. ponticum* and *T. timopheevi*, the majority of introgression should exist in smaller chromosome size in our panel. To address the limitations of a single reference genome and improve the detection of alien gene expression, we constructed a pan-gene atlas from 24 published *Triticeae* genomes. If we simply use a combination of different genomes as reference when doing the RNA-seq reads mapping, gene sequences conserved across species will not be properly handled due to the limitation of short reads alignment algorithms. Thus, we developed a workflow to construct a non-redundant pan-gene atlas.

First, all 107,892 genes from the Chinese Spring (CS) reference genome were retained (Fig. 2d). Second, to study the expression patterns of large introgressed chromosomal fragments, 5,602 genes from four large introgressions in our wheat panel were included (Fig. 2d). Third, to maximize the study of small introgressed genes not present in CS, non-redundant genes from 19 *Triticeae* genomes with different ploidy levels were selected using the following two steps:1) Genes homologous to those in the Chinese Spring genome were removed. We used OrthoFinder to classify 132,527 ortholog groups from the 20 genomes based on homology, resulting in orthogroups containing 1 to 4,871 genes (Fig. 2a, b). The largest gene family is linked to ribonuclease H-like, zinc finger (CCHC-type), and DNA/RNA polymerase domains. A total of 44,798 ortholog groups containing at least one CS gene were excluded, and 87,729 groups, containing genes from up to 19 other genomes, were retained (Fig. 2c), 2) for the 87,729 orthologous groups, the longest transcript was retained. Variable genes (42,370) were shared by more than two genomes, and private genes (45,359) were found in only one genome (Fig. 2d). The integration of the three gene sets led to the creation of a non-redundant pan-gene atlas consisting of 201,223 genes (Fig. 2d). This valuable resource offers high-quality gene expression quantification and enables the study of alien fragment functions, helping to explore how germplasm diversity affects gene expression diversity.

**Fig. 2.**
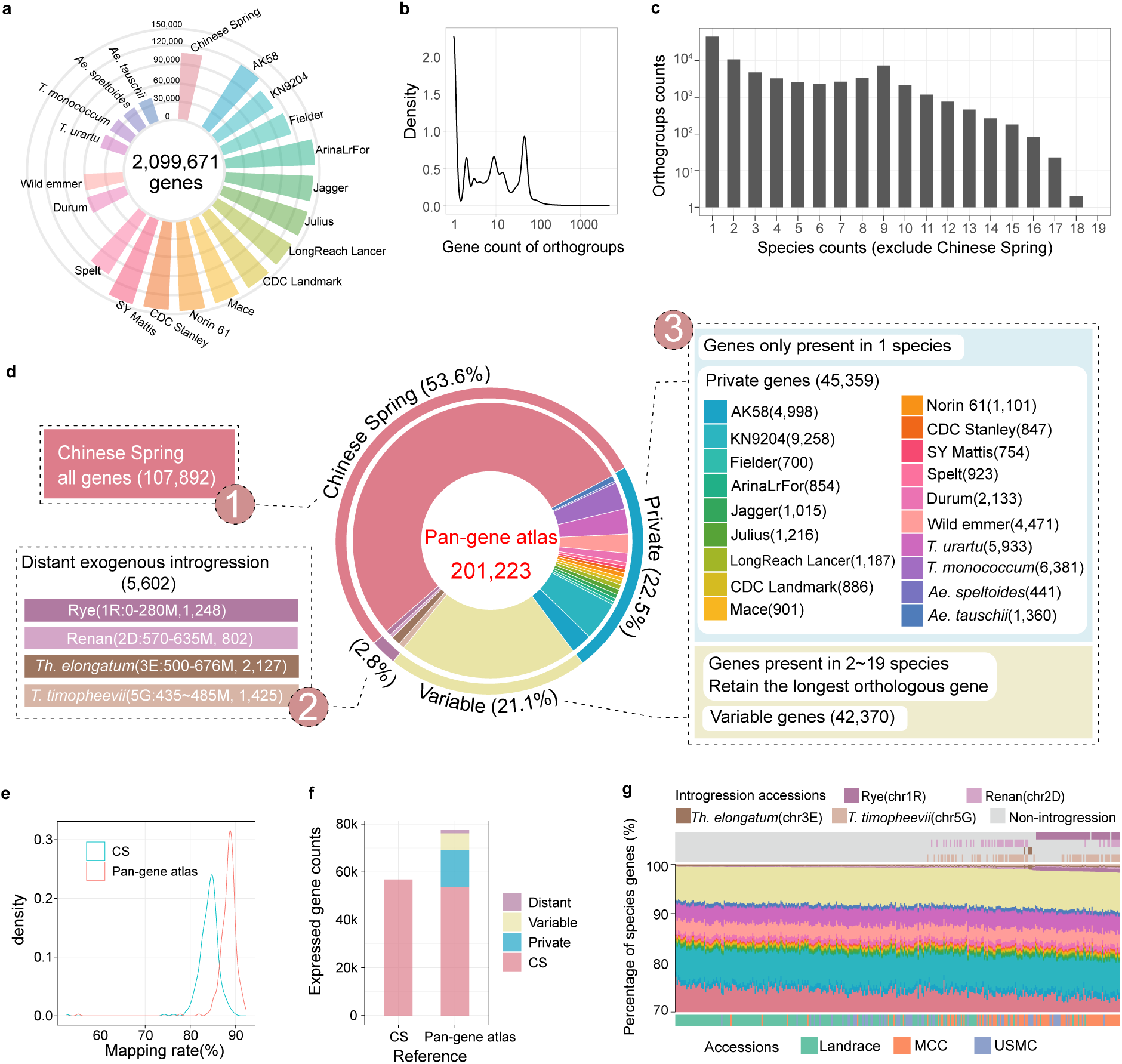
Creating a pan-gene atlas. **a** The number of high-confidence genes in 20 genomes of wheat varieties and relative species at different ploidy levels. **b** Density distribution of the number of homologous genes in each orthologous group. **c** The number of genomes contained in each orthologous group (excluding groups containing Chinese Spring genes). **d** Construction of the pan-gene atlas from three parts: (1) all high-confidence genes of Chinese Spring, (2) genes from large introgressed fragments detected for our panel, and (3) variable genes and private genes from the 19 remaining genomes. **e** Density distribution of the mapping rates for RNA-seq data from 328 wheat accessions aligned separately to Chinese Spring and the pan-gene atlas as reference gene models. **f** The number of expressed genes for 328 wheat accessions, separately quantified using Chinese Spring and the pan-gene atlas as reference. **g** Heatmap showing the proportion of expressed genes for each wheat accession. The bottom color bar indicates the type of wheat accessions, the middle heatmap shows the proportion of expressed genes from different genomes (genome colors are consistent with Fig. 2d), and the top heatmap represents the introgression of foreign fragments in different wheat varieties.

To evaluate the alignment accuracy and gene detection improvement, RNA-seq data from all wheat samples were mapped to the pan-gene atlas. The average alignment rate of the pan-gene atlas was 88.4%, representing a 4.3% improvement compared to the single CS reference genome (Fig. 2e and Supplemental Table 3). The pan-gene atlas detected 77,447 genes with TPM > 0.5 in at least 5% of samples, including 53,645 (69.3%) CS genes, 7,036 (9.1%) variable genes, 15,489 (20.0%) private genes, and 1,277 (1.6%) distant genes, detecting 20,615 more genes than the single CS reference (Fig. 2f). Expressed CS genes accounted for 71.88-77.19% in per sample, variable genes for 6.66-8.28%, private genes for 15.65-18.69%, and distant genes for 0.42-1.70%. And 53 samples contained more than one introgression segment (Fig. 2g and Supplemental Table 4, 5). Interestingly, MCC and USMC exhibited a significantly higher number of expressed genes than landraces, particularly for non-CS genes (Supplementary Fig. 4, 5), suggesting that modern breeding has improved gene expression diversity in cultivars.

### The gene expression pattern of introgression segments from rye, *Ae. markgrafii*, *T*. *ponticum* and *T*. *timophevii*

Wild relative introgression segments with novel genes or regulatory elements can interact with the wheat genome, altering expression patterns and agronomic traits. Recent studies indicate that species-specific genes carried by introgression segments are rarely expressed, while those replacing wheat homologs tend to be downregulated [25]. To uncover the changes in the expression patterns of wild relative introgression segments from donors to wheat introgression lines, we mapped RNA-seq data from 10 rye samples, 2 Renan samples (excluded from the main text due to small sample size; see supplementary materials), 7 *T*. *ponticum* samples, and 3 *T*. *timophevii* samples to a pan-gene atlas combined with the donor genome (Supplemental Table 6). We analyzed the expression changes of 5,602 distant alien genes between donor samples and wheat introgression lines. The proportion of expressed genes within introgression segments was significantly lower in wheat than in their donors (Fig. 3a, f and Supplementary Fig. 6d, e). Overall, both species-specific and conserved introgressed genes showed suppressed expression in the wheat background compared to that in the donor background (Fig. 3b, g and Supplementary Fig. 6a, f, g). The conservation of a gene is positively correlated with the likelihood of being expressed (Fig. 3c, h and Supplementary Fig. 6h, i), which is consistent with earlier studies in model organisms [32–34].

**Fig. 3.**
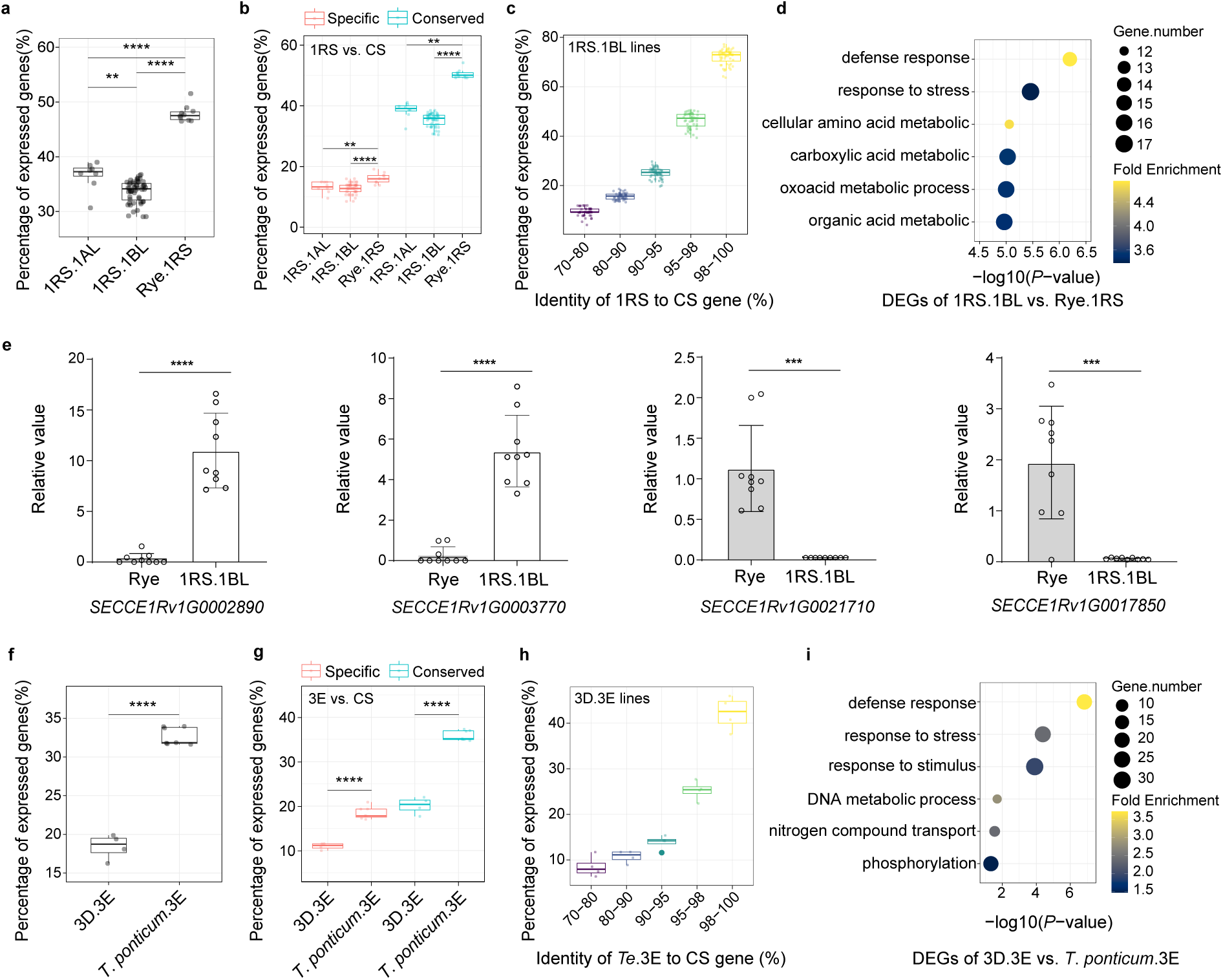
Expression pattern of distant introgression in wheat and their donors. **a** The percentage of expressed genes within the chromosome 1RS (0-280 Mb) region of rye translocation lines 1RS.1AL, 1RS.1BL and rye lines. **b** The percentage of expressed specific and conserved genes within the chromosome 1RS (0-280 Mb) region in rye translocation lines 1RS.1AL, 1RS.1BL, and rye lines. **c** The relationship between cross-species gene sequence conservation and proportion of expressed genes. **d** Functional enrichment analysis of differentially expressed introgressed genes between the 1RS.1BL translocation lines and rye lines. **e** Transcript-level validation of differentially expressed genes (DEGs) between the 1RS.1BL translocation lines and rye lines using qRT-PCR, with *Tatublin* as the internal control. \**p* < 0.05, \*\**p* < 0.01, \*\*\**p* < 0.001, \*\*\*\**p* < 0.0001 (two-tailed Student’s *t*-test). **f-i** Analytical figures of *T*. *ponticum* introgression lines using *Th*. *elongatum* as the reference, corresponding to the analyses in panels (a-d).

In 1RS.1BL translocation lines, 138 genes were upregulated and 223 downregulated, while the *T*. *ponticum* introgression lines had 289 upregulated and 247 downregulated genes. Upregulated genes in both cases were enriched in disease resistance and environmental adaptation, while downregulated genes were linked to fundamental biological functions (Fig. 3d, i). The 1RS.1BL and *T*. *ponticum* introgression segments are well-known for their disease resistance, such as *Lr10* (ortholog: *SECCE1Rv1G0000670*) for leaf rust resistance and *Pm8* (ortholog: *SECCE1Rv1G0001880*) for powdery mildew resistance [35],which were identified as upregulated in 1RS.1BL translocation lines. The resistance genes *Lr24* (leaf rust) and *Sr24* (stem rust), derived from *T*. *ponticum* [36] were also upregulated in introgression. These directional changes of gene expression activity are likely a result of breeding selection since wild relatives have been continuously utilized to improve the disease resistance of wheat during the last century. *T*. *timophevii* has also provided resistance genes for wheat breeding [37]. We did not observe GeneOntology enrichment for disease resistance among DEGs (Supplementary Fig. 6j), probably because the number of genes on this introgression were too small to statistical tests.

To verify the enriched differential genes in the introgression lines, we selected three upregulated NLR genes and three downregulated core metabolic genes in 1RS.1BL for genomic DNA amplification to check gene presence/absence. These three NLR genes were not present in all rye lines but were found in the 1RS.1BL introgression lines, which may be one of the important reasons why NLR genes in introgression lines exhibit upregulation at the population level, while the core metabolic genes were present in all samples, suggesting these genes were not present in the original rye used for distant hybrid breeding (Supplementary Fig. 6b). qRT-PCR further confirmed the gene expression changes, displaying consistent results with differential expression analysis based on short reads sequencing (Fig. 3e and Supplementary Fig. 6c; Supplemental Table 7,8). Those regulatory changes are consistent among rye, *T*. *ponticum* and *T*. *timopheevi* suggesting they might be due to continuous breeding selection. Although introgression from wild relatives is a common practice in wheat breeding, our results showed that not every introgressed gene are equally activated. Among the set of traits contributed by wild relatives, disease resistance and tolerance to environmental stress are the most famous [38, 39]. Perhaps it is not surprising genes involved in basic cellular process were suppressed, since such genes have already existed in the wheat genome. We next investigated the how those introgressed genes are regulated by the wheat genome using our population RNA-seq data.

### eQTL map of pan-gene atlas

To discover genetic loci regulating gene expression, we conducted association analysis between the expression levels of each gene and genome-wide SNPs based on CS reference. This analysis identified eQTLs that exhibit significant associations with variation of expression levels. To better analyze the regulatory relationship between variation sites and gene expression, the eQTLs can be classified into three categories: intergenic-eQTLs, which are regulatory loci found in intergenic regions; inactive-eQTGs, which are located on genes that are not expressed in the population; and active-eQTGs, which are located on genes that are expressed in the population. Genes regulated by eQTLs are called eGenes (Fig 4a). A total of 38,409 eGenes were identified, including 25,176 (65.5%) eGenes from Chinese Spring, 8,016 (20.9%) private eGenes, 4,157 (10.8%) variable eGenes, and 1,060 (2.8%) distant eGenes (Fig. 4b). At the population level, 39,038 (51%) non-eGenes had no regulatory genetic loci (Supplementary Fig. 7a, b), suggesting their expression variation might be predominantly influenced by environmental factors. 32,264 (84.0%) eGenes were regulated by intergenic-eQTLs, inactive-eQTGs, active-eQTGs, while 172 (0.4%) eGenes were exclusively regulated by active-eQTG and 117 (0.3%) by inactive-eQTG. Notably, 3,414 (8.9%) eGenes were only regulated by intergenic-eQTLs, highlighting the importance of intergenic regions in gene regulation (Fig 4c).

**Fig. 4.**
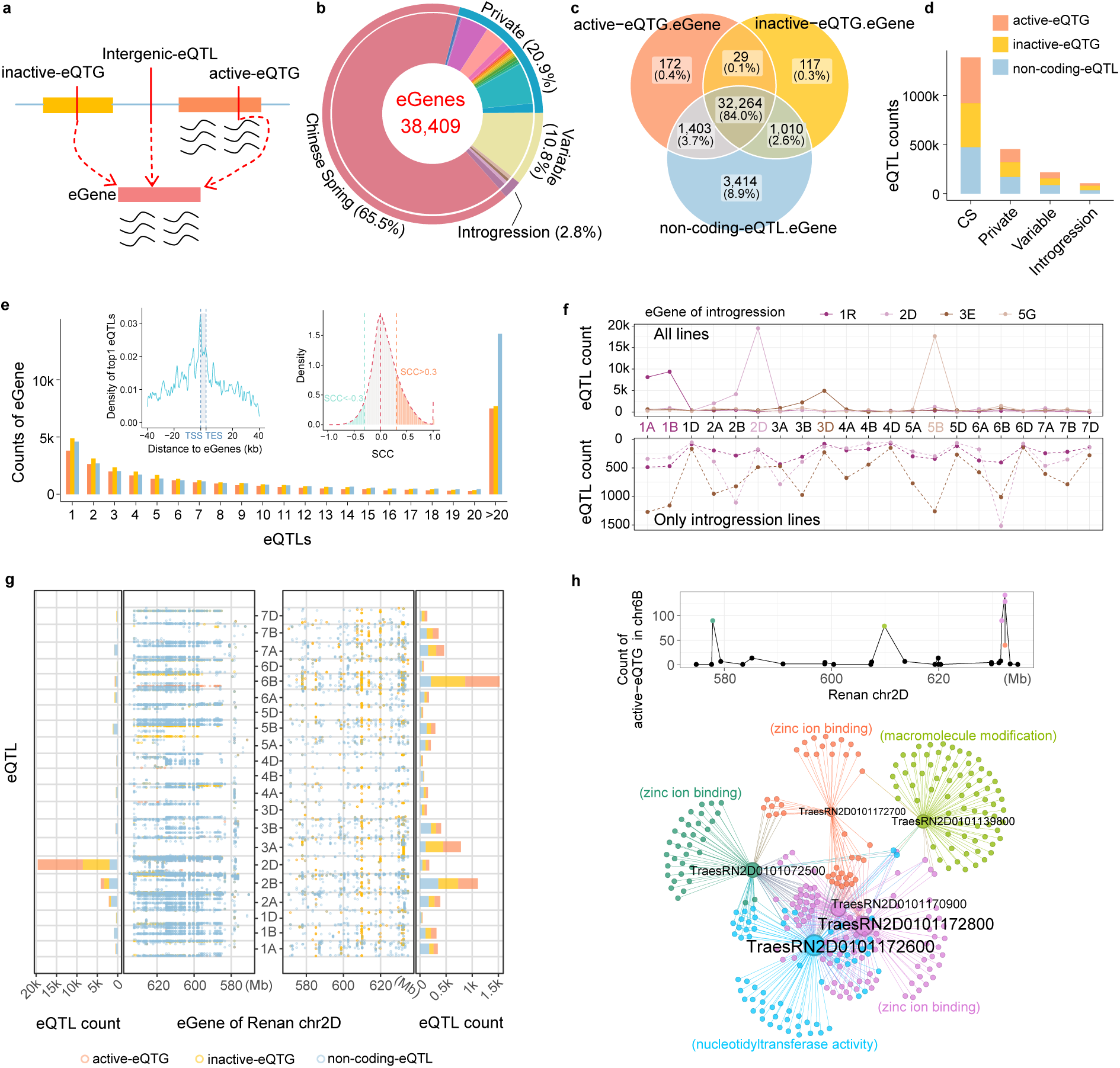
eQTL map of the pan-gene atlas. **a** A schematic model of eQTLs regulating eGenes. Active-eQTG indicates that the regulatory locus is located within a gene and expressed in the population, whereas inactive-eQTG indicates that the locus is within a gene but not expressed in the population. Intergenic-eQTL refers to regulatory loci located in intergenic regions. **b** The distribution of gene types and the number of eGenes. The middle color bar represents genome classification, with colors consistent with Fig. 2d. **c** A venn diagram showing the distribution of eGenes regulated by three types of eQTLs. **d** The number of eQTLs for eGenes in the pan-gene atlas. **e** The outer figure shows the distribution of eQTLs corresponding to each eGene. The inner left panel shows the physical distance distribution between the most significant eQTLs and their eGenes (positive values indicate downstream of eGenes, negative values indicate upstream). The inner right panel illustrates the Spearman’s rank correlation coefficient (SCC) distribution between active-eQTGs and their eGenes. **f** The number of eQTLs for distant introgressed genes calculated for all samples and translocation lines, with the number of eQTLs on each chromosome displayed. **g** The eQTL map of introgressed genes on Renan chr2D. The Y-axis represents the position of SNP on wheat chromosomes, the dot plot X-axis represents the position of introgressed eGenes on Renan chr2D, and the bar plot X-axis represents the number of eQTLs. **h** The bar chart on the right panel (g) shows the regulatory relationships of active-eQTGs on wheat chromosome 6B with eGenes on Renan chr2D. In the line chart above, the X-axis represents the position of eGenes on Renan chr2D, and the Y-axis represents the number of active-eQTGs on wheat chromosome 6B. Different colored points correspond to different eGenes in the regulatory network diagram below. The regulatory network diagram shows the relationships between the eGenes (colored points in the line chart above) and their corresponding active-eQTGs on chromosome 6B.

eGenes were regulated by 2,168,336 eQTLs (Fig 4d). The number of eGenes regulated by a single eQTL is the highest, and the number of eGenes regulated by multiple eQTLs decreases as the number of eQTLs increases (Fig. 4e). The physical distance between eGenes and eQTLs may influence the strength of regulation [40]. We observed that the strongest regulatory signals are concentrated near gene regions, particularly around the transcription start site (Fig. 4e). So, the syntenic chromosome location of the non-CS genes can be anchored by the strongest eQTL signal located at the Chinese Spring genome. In this study, eQTLs located on the same chromosome as the eGene in CS were classified as *cis*-eQTL, while those on different chromosomes were classified as *trans*-eQTL. The significance of *cis*-eQTL signal is much stronger than *trans*-eQTL of the same gene (Supplementary Fig. 7c, d).

Furthermore, there are 631,512 pairs of active-eQTGs and eGenes with an absolute Spearman correlation coefficient greater than 0.3 (|SCC|>0.3), of which 489,805 pairs were positively correlated and 141,707 pairs were negatively correlated (Fig. 4e). In addition, 91% of intergenic-eQTLs are located in transposable element (TE) regions suggesting TE play a role in gene regulation and potentially impacts the agronomical traits. The proportion of TE types for intergenic-eQTLs are comparable to those reported in previously published results in wheat [41], suggesting a uniform dispersion of these intergenic-eQTLs throughout TE regions (Supplementary Fig. 7e).

### *Trans*-regulation of introgressed genes from rye, *Ae. markgrafii*, *T*. *ponticum* and *T*. *timophevii*

To analyze the regulation of introgressed genes by wheat, we calculated eQTLs for translocation lines of 1RS.1BL, *Ae*. *markgrafii*, *T*. *timophevii*, separately. eQTL results of distant introgressed genes from 328 samples at the population level were used for comparison (Supplemental Note). The number of eGenes located on 1RS across all lines was 445, whereas only in 1RS.1BL introgression lines, with only 257 eGenes identified (Supplementary Fig. 8a). The number of eQTLs calculated using only introgression lines was significantly lower compared to those identified across all lines (Supplementary Fig. 8b). The eQTLs of distant introgressed genes are concentrated on chromosomes 1A, 1B, 2D, 3D, and 5B (Fig 4f, g and Supplementary Fig. 8c, e, f), which is influenced by the population structure. These eQTLs mainly reflected the presence/absence variation of introgressed segments, rather than the regulatory mechanisms by which wheat regulated introgressed genes. In contrast, eQTLs calculated from introgression line samples are more evenly distributed (Fig 4f, g and Supplementary Fig. 8c, 8e, 8f), better capturing how wheat genome regulates introgressed genes.

The 2D genes introgressed from *Ae. markgrafii* have the highest number of eQTLs on chromosome 6B (Fig. 4f). We found 6 genes with 1,168 eQTLs, including 570 active-eQTGs (Fig. 4g). The regulatory network of active-eQTGs and eGenes is associated with zinc ion binding (Fig. 4h). Zinc binding plays key roles in plant growth, development, metabolism, stress tolerance, and immunity. Zinc deficiency causes stunted growth and chlorosis, making zinc regulation and supplementation crucial for plant health [42, 43]. Additionally, the active-eQTG *TrarsCS5A02G120000* was a regulatory hotspot for 1RS genes among the distant introgressed genes (Supplementary Fig. 8c), and it regulated 94 eGenes of all 1RS.1BL lines (Supplementary Fig. 8c). This active-eQTG is located in the centromeric region of chromosome 5A. The centromeric region of chromosome 5A in 1RS.1AL and 1RS.1BL has undergone variation compared to the reference genome. When this region undergoes genomic sequence variation, the samples are introgression lines (Supplementary Fig. 8d), suggesting that the centromeric structure of chromosome 5A may influence the ability of integrating external chromosomes. The regulation of introgressed genes involves multiple chromosomes, and this cross-chromosomal regulatory feature indicates that wheat exhibits a complex genetic regulatory network when adapting to different foreign genes. These conclusions provide a theoretical foundation and practical reference for future research on wheat genetic improvement, the adaptability of introgressed genes, and the mechanisms of stress resistance.

### Multi-omics modeling identified candidate genes from the pan-gene atlas for agronomic traits

Recent studies have shown that both transcriptome-wide association study (TWAS) and summary data-based Mendelian randomization (SMR) integrate GWAS and eQTL data to identify genes associated with complex traits: TWAS employs gene expression prediction and transcriptome-wide association analysis to uncover associations between *cis*-regulated gene expression and traits, while SMR utilizes Mendelian randomization to explore causal relationships between gene expression and traits [44, 45]. Therefore, we used the eQTL map of the pan-gene atlas to integrate GWAS, TWAS, SMR and spearman correlation coefficient (SCC) between expression levels and phenotypic values, enabling the identification of CS and non-CS candidate genes that affect phenotypic variation (Fig. 5a). For our panel, we collected 34 phenotypes for field agronomic traits and infection phenotypes of 8 powdery mildew isolates (Supplemental Table 9). The genes detected by only a single method were removed from further analysis. A total of 209 candidate genes were obtained for 34 field agronomic traits with 51 non-CS genes (Fig. 5b, c and Supplementary Fig. 9,10; Supplemental Table 10, 11, 12, 13, 14). Given that most of cloned powdery mildew resistance genes are NLRs or kinases [46], only 22 genes of those types were treated as candidates (Fig. 5b and Supplementary Fig. 11; Supplementary Table 15, 16, 17, 18, 19), including 3 non-CS genes (Fig. 5c).

**Fig. 5.**
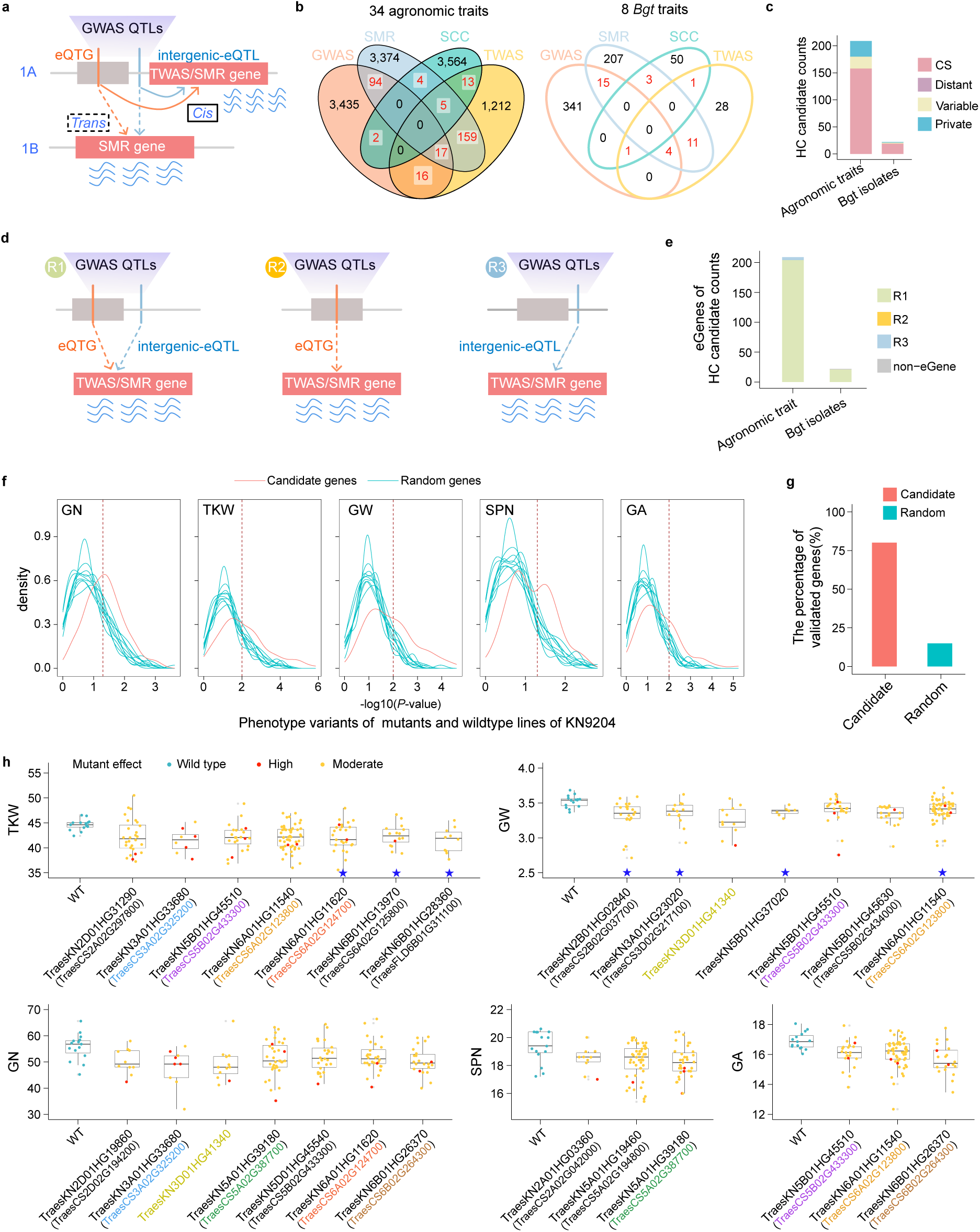
Joint eQTLs of the pan-gene atlas and GWAS, TWAS, SMR analysis of 34 agronomic traits and 8 *Bgt* isolates infection phenotypes in wheat. **a** Schematic diagram of the predicted candidate gene model. The GWAS QTL candidate regions include signals from eQTG and intergenic-eQTL. Candidate genes are predicted using *cis*-eQTLs through TWAS, and both *cis-* and *trans-*eQTLs through SMR. **b** Venn diagram of the distribution of candidate genes predicted by GWAS, TWAS, SMR, phenotypes, and gene expression SCC. **c** The number and proportion of candidate genes classified by the pan-gene atlas. **d** Three eQTL regulatory patterns assist in identifying candidate genes. R1 represents candidate genes regulated by both gene and intergenic regions, R2 represents candidate genes regulated only by the gene region, and R3 represents candidate genes regulated only by the intergenic region. **e** The types and numbers of regulatory patterns for the candidate genes. **f** Verification of candidate genes using the indexed Kenong9204 (KN9204) mutant library. The phenotypes of mutants and wild-type for the predicted candidate genes were analyzed using a two-tailed Student’s *t*-test, with the red line representing the *t*-test distribution for the candidate genes. Randomly selected genes in 10 times, with each selection containing the same number of random genes as the candidate genes. Blue-green line representing the *t*-test p-value distribution for the randomly selected genes. **g** The proportion of verified candidate genes and random genes in the KN9204 mutant library. **h** Representative candidate genes verified using the KN9204 mutant library. Blue pentagrams indicate that the gene was predicted to be associated with TKW or GW and the corresponding phenotype was validated in mutants. The remaining 20 candidate genes were predicted to be associated with multiple agronomic traits and validated in relevant traits (TKW, GW, GN, SPN, GA) in the mutant library. Red points represent premature termination (high effect) and yellow points represent non-synonymous mutations (moderate effect) in the candidate gene. Blue-green points represent wild-type.

GWAS can only identify significant SNPs, whereas TWAS and SMR directly identify candidate genes [47, 48]. For candidate genes with an eQTL, there are three regulatory models. 99% of the candidates are eGenes, with 95%-98% regulated by both genic and intergenic regions. 2%-5% are regulated only by intergenic regions, highlighting the critical role of non-genic regions (Fig. 5d). *Ppd*-*D1* is regulated by the R1 model. *Ppd*-*D1*, previously reported to be involved in photoperiod response [17], was identified as a candidate gene for life cycle regulation in our TWAS analysis. A frameshift mutation (CCGACG→C) in the *Ppd*-*D1* gene led to a significant change in expression levels, ultimately affecting the lifecycle (Supplementary Fig. 12a). Using the SMR approach, we identified the gene *TaGL3*-*5B*, associated with grain length and width. *TaGL3-5B* (Supplementary Fig. 12b), which is involved in grain size [49], is regulated by the R3 model and only controlled by intergenic-eQTLs (Supplementary Fig. 12b). We also identified the cloned *Pm4* allele [50] (*GWHGANRF020664* from Aikang58) and *Pm3* (*TraesCS1A02G008100*) [51] using the SMR approach, the *Lr10* allele [35] (*SECCE1Rv1G0000670* from rye) using TWAS, and the *Pm5* alleles (*TraesCS7B02G441700*) [52] using GWAS. Additionally, the candidate *TraesCS3D02G201900* of biomass per plant at jointing stage has no SNP variation in the population but is regulated by *cis*-eQTL. TWAS and SMR associate this gene with phenotypic variation, suggesting that eQTL may indirectly affect phenotypic variation by altering gene expression levels. Identification of cloned genes not encoded by the Chinese Spring reference genome demonstrated the power of using pan-gene atlas as reference when mapping RNA-seq short reads.

To validate the reliability of candidate genes from the pan-gene atlas, we utilized the indexed EMS library of the wheat cultivar Kenong9204 (KN9204) [53]. For the 99 agronomic candidate genes, either directly from the KN9204 genome or with a KN9204 homolog, we queried their mutant lines from the approximately 2000 EMS lines in the library. Among these, 72 candidates have at least 5 mutant lines carrying either knockout or non-synonymous mutations. We collected seven traits including GA, GL, GP, GW, SPN, TKW and GN for mutant lines as well as wild type cultivars. Significant phenotypic difference was observed for 58 (80.6%) out of 72 candidates, including 14 non-CS genes among the validated ones. As a comparison, only 14.9% of randomly selected genes demonstrated this level of phenotypic changes, highlighting the high reliability of our identified candidate genes (Fig. 5f, 5g and Supplementary Fig. 13a, 13b; Supplementary Table 20; Supplementary Note). The *TraesKN3D01HG41340* mutant significantly reduced grain width and spike grain number, with the strongest effect from premature termination mutations. For the grain width candidate gene *TraesKN5B01HG37020*, its mutant showed a significant reduction in grain width. The remaining candidate genes are homologs of KN9204, among which 14 validated KN9204 genes contain premature termination mutations, leading to significant phenotypic variation (Fig. 5h and Supplementary Table 21). In summary, we provided a reliable method for identifying exogenous candidate genes and validated them using a mutant library, offering gene resources for practical applications in production.

### Differentially expressed genes and modules between different sub-populations

In order to understand the impact of breeding effort on gene expression, we wondered what genes are highly expressed or suppressed in individual sub-populations or breeding programs. We are also wondering whether differentially expressed genes (DEGs) are involved in trait improvement. First of all, we observed dramatic expression changes between sub-populations consistent with the genetic divergence. There were 7,413 DEGs between MCC and LR, including 2,668 (36.0%) non-CS DEGs; 7,517 DEGs between USMC and LR, including 2,312 (30.8%) non-CS DEGs; and 3,813 DEGs between USMC and MCC, including 1,755 (46.1%) non-CS DEGs (Fig. 6a). This indicated that the differences between breeding programs are smaller than those between landraces and cultivars. The 30∼46% DEG being non-CS genes highlighted the limitations of using a single reference genome in RNA-seq studies.

**Fig. 6.**
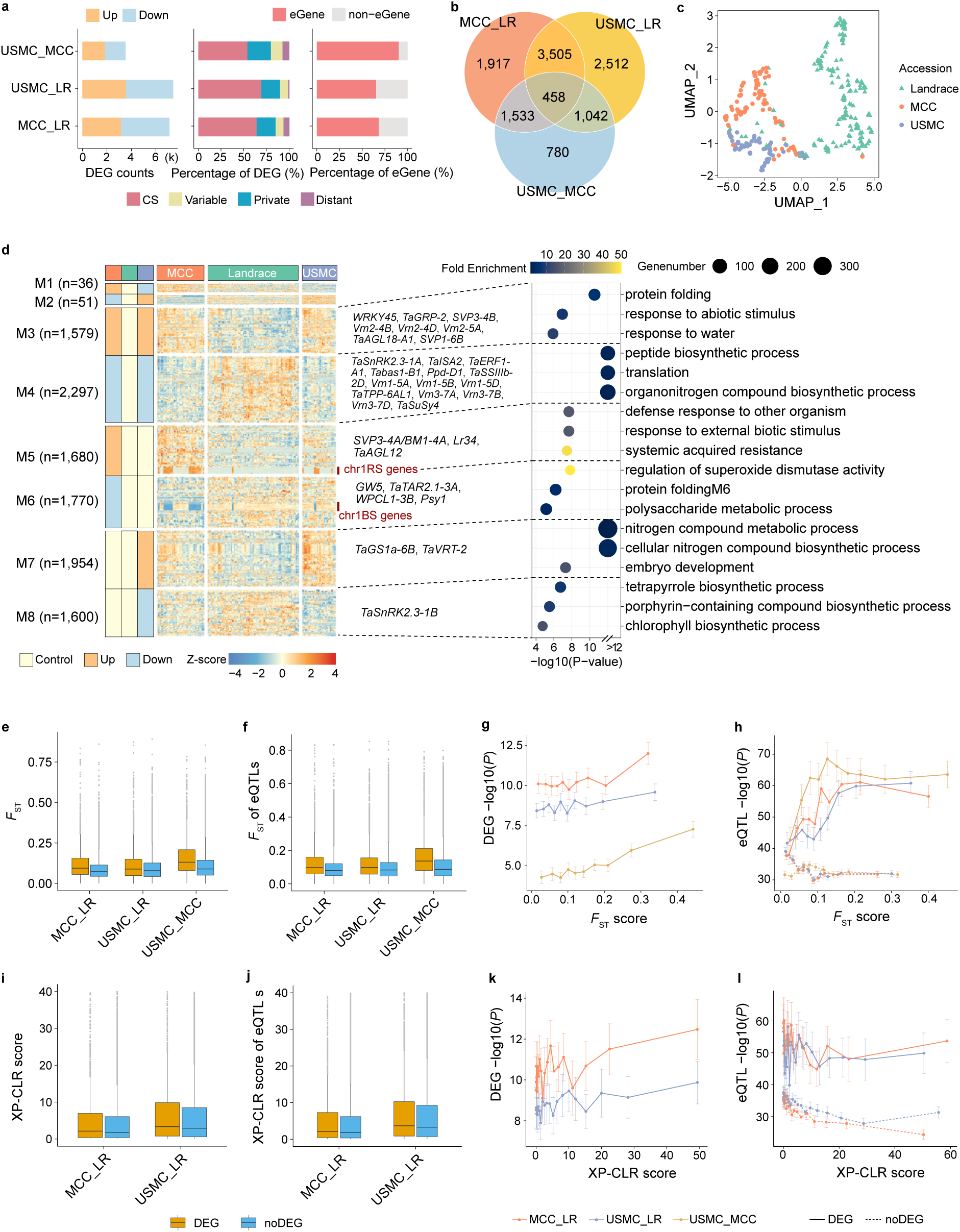
The genome-wide impact of breeding selection on gene expression regulation. **a** The number of DEGs for each pair of sub-populations. **b** Sharing of DEGs between sub-populations. **c** UMAP plot of the dimensionality reduction analysis of DEGs expression matrix. **d** Heatmap of the 8 modes of DEGs. The numbers on the left represent the count of DEGs. The dashed box in the middle highlights representative cloned genes of known function among the DEGs, and the right side shows functional enrichment for each module (M3-M8). **e** *F*_ST_ of the genic region for DEGs. **F** *F*_ST_ of eQTLs for DEGs. **g** The relationship between the *F*_ST_ values of DEGs and the strength of differential expression. **h** The relationship between the *F*_ST_ values of eQTLs for DEGs and the strength of eQTL signals. **i** XP-CLR score of DEGs. **j** XP-CLR score of eQTLs for DEGs. **k** The relationship between the XP-CLR scores of DEGs and the strength of differential expression. **l** The relationship between the XP-CLR scores of eQTLs for DEGs and the eQTL signals.

Secondly, we investigated if DEGs were genetically regulated by SNP. More than 65% of DEGs between cultivar and landraces have at least one eQTL, which is significantly over-represented compared to the genomic background (49%) (Fisher’s exact test, *p*-value=0.0319) (Fig. 6a). Nearly 90% of the DEGs between MCC and USMC have eQTLs, suggesting that the gene expression differences between those two counties were primarily driven by genetic effects. The fact that DEGs between sub-populations tend to be genetically controlled indicated the role of breeding selection in shaping the gene expression landscape. UMAP-based dimensionality reduction analysis of the expression of these DEGs clearly distinguished the sub-populations, indicating that the genetic divergence is coupled with transcriptomic differentiation (Fig. 6b, c).

To investigate the similarities and differences in breeding directions between MCC and USMC, DEGs were categorized into eight modules (Fig. 6d, Supplementary Fig. 12). We found that genes upregulated in both MCC and USMC in module M3 during the seedling stage were enriched in functions related to abiotic stimulus and response to water. This indicates that both MCC and USMC improved the expression of genes associated with external stimuli and water response during the seedling growth phase compared to LR. Genes downregulated in both MCC and USMC in module M4 were enriched in functions suggesting reduced protein synthesis, translation efficiency, and nitrogen synthesis during the seedling stage. This could be one of the reasons why nitrogen fertilizer supplementation is necessary during this growth stage. Notably, module M5 consisted of genes specifically upregulated in MCC, which were significantly enriched in resistance-related responses. This indicates that, in addition to the resistance genes commonly selected in M3, Chinese breeders selected more resistance-related genes. Conversely, module M7 consisted of genes specifically upregulated in USMC, which were significantly enriched in nitrogen compound biosynthesis, compensating for the reduction in nitrogen efficiency observed in M4 (Fig. 6d). These findings probably indicate that while both China and the United States focused on improving wheat resistance and nitrogen efficiency, Chinese breeders placed greater emphasis on resistance breeding, whereas U.S. breeders concentrated more on enhancing nitrogen utilization efficiency.

The cloned DEGs provide a more detailed reflection of the similarities and differences in breeding selection between China and the United States. For example, Module M3 includes *TaGRP-2*, *TaAGL18, VRN2*, *SVP3, WRKY45* and MADS-box transcription factor family [54–56], which are linked to environmental stress responses and flowering time regulation. Module M4 includes *TaISA2*, *Tabas1*, *TaSSIIIb*, *TaTPP*-*6AL1*, and *TaSuSy4*, the genes downregulated in MCC and USMC are enriched in peptide biosynthesis, protein translation, and the biosynthesis of organic nitrogen compounds [6, 57–60]. Additionally, genes involved in photoperiod and flowering also show up in this module, including *Ppd-D1*, *VRN1* and *VRN3*. (Fig. 6d). Compared to landraces, *VRN2* in module M3 is upregulated in both MCC and USMC, while *VRN1* and *VRN3* in module M4 are downregulated. In winter wheat, vernalization ensures that flowering does not occur prematurely by maintaining low *VRN1* expression and high *VRN2* expression, which inhibits *VRN3* until low temperatures trigger *VRN1* induction [61]. In this study, spring wheat varieties were more prevalent in landraces, while winter wheat varieties were more common in cultivars (Supplementary Fig. 14). This finding was also validated using 1,034 accessions from previously published datasets (Supplementary Fig. 15) [5]. In modern breeding, winter wheat has been more frequently selected due to its ability to overwinter and store more nutrients, resulting in higher yields [62, 63].

In module M5, a greater abundance of 1RS gene expression was detected in MCC, and the 1RS gene was also present in some varieties of USMC, indicating the widespread application of rye genes in modern cultivars. *Lr34* is upregulated in MCC, indicating that MCC has a higher frequency of activated *Lr34* [64]. The breeding application of *Lr34*/*Yr18*/*Sr57*/*Pm38*/*Ltn1*/*Sb1* has a history of over 100 years, and it continues to provide robust resistance against several wheat diseases [65]. In M6, genes related to yield and quality, such as *GW5*, *TaTAR2.1-3A*, *WPCL1-3B*, and *Psy1*, were downregulated at the seedling stage compared to landrace, which may indicate that their expression increases at later stages [66–69]. In M7, *TaGS1a-6B*, which encodes *Glutamine Synthetase 1* (*GS1*), a key enzyme in plant nitrogen metabolism [70, 71]. *GS1* catalyzes the reaction that synthesizes glutamine from glutamate and ammonium ions, and the expression level of this gene is closely associated with nitrogen absorption and utilization efficiency in wheat [71]. The utilization of this gene in America may be related to the characteristics of low planting density and reduced application of chemical fertilizers (Fig. 6d). In summary, MCC and USMC have applied more resistance genes in the breeding improvement process, with MCC focusing more on resistance breeding and USMC placing more emphasis on enhancing nitrogen efficiency.

### Population divergence and selective scan of differentially expressed genes

To investigate genetic differentiation of DEGs in modern breeding, we performed an *F*_ST_ analysis between DEGs and non-DEGs. The *F*_ST_ of the genomic regions containing DEGs were significantly higher than the *F*_ST_ of the regions harboring non-DEGs (Wilcoxon test, *p*-value < 2.2e-16 for MCC vs. LR, USMC vs. LR, and USMC vs. MCC). Additionally, the *F*_ST_ of the eQTL regulating DEGs were also significantly higher than that of the eQTLs associated with non-DEGs (Wilcoxon test, *p*-value < 2.2e-16 for MCC vs. LR, USMC vs. LR, and USMC vs. MCC). This result indicates that not only were the genetic differentiation of DEGs higher, but the genetic differentiation of the eQTLs regulating these DEGs were also higher (Fig. 6e, f). Furthermore, as *F*_ST_ increases, the signals of the DEGs or their eQTLs were also stronger, while the signals of the eQTLs regulating non-DEGs were significantly lower (Fig. 6g, h).

To further investigate whether DEGs were under to natural selection, we performed maximum likelihood ratio XP-CLR analysis for cultivars compared to landraces. The XP-CLR scores for DEGs in both MCC and LR, as well as USMC and LR, were significantly higher than non-DEGs (Wilcoxon test, *p*-value = 7.75e-10 for MCC vs. LR, and *p*-value = 1.649e-05 for USMC vs. LR). Additionally, the XP-CLR scores of the eQTL loci regulating DEGs were also significantly greater than those of the eQTLs for non-DEGs (Wilcoxon test, *p*-value = 1.769e-10 for MCC vs. LR, and *p*-value = 3.352e-07 for USMC vs. LR), indicating that DEGs have been selected during the breeding process (Fig 6i, j). Moreover, as the XP-CLR scores of the genomic regions containing DEGs increase, the signals of the DEGs become stronger; although there was a slight upward trend in the signals of eQTLs regulating DEGs with increasing XP-CLR scores, the signals of eQTLs for non-DEGs showed a slight downward trend (Fig 6k, l). Overall, DEGs and their associated regulatory loci have displayed significant genetic differentiations and have also been influenced by stronger selection pressure.

### Modern breeding reshaped the gene regulatory network of wheat

To investigate changes in gene co-expression and regulatory networks during modern breeding, this study utilizes an effective approach by analyzing shifts in co-expression patterns. Specifically, the number of correlations between active-eQTGs and eGenes with |SCC| > 0.3 is calculated across different sub-populations (see “Methods”). A total of 1,046 DEGs from LR and MCC, and 616 DEGs from LR and USMC, are not only regulated by active-eQTGs but are also identified as candidate genes for agronomic traits by at least one method (Fig. 7a). The 1,046 candidate genes were regulated by 50,010 active-eQTGs, with 27,157 pairs (54.3%) and 43,786 pairs (87.6%) of active-eQTGs and eGenes showing co-expression (|SCC| > 0.3) in the landrace group and MCC group, respectively (Supplementary Table 22, 23). The co-expression proportion in the MCC group was significantly higher than in the landrace group (Fisher’s exact test, *p*-value < 2.2e-16) (Fig. 7b). Similarly, the 616 candidate genes were regulated by 15,123 active-eQTGs, with 8,565 pairs (56.6%) and 12,825 pairs (84.8%) of active-eQTGs and eGenes showing co-expression in the landrace group and USMC group, respectively (Supplementary Table 24). The co-expression proportion in the USMC group was significantly higher than in the landrace group (Fisher’s exact test, *p*-value < 2.2e-16) (Fig. 7b). *SVP1*-*6B*/*BM10*-*6B* was a cloned gene associated with wheat spike and spikelet development, flowering, and plant height [72]. Its co-expression regulatory relationships were fewer in landraces compared to MCC and USMC (Fig. 7c). This indicated that modern breeding, transitioning from landraces to MCC or USMC, has led to increasingly complex and highly connected co-expression regulatory networks of phenotype-associated genes.

**Fig. 7.**
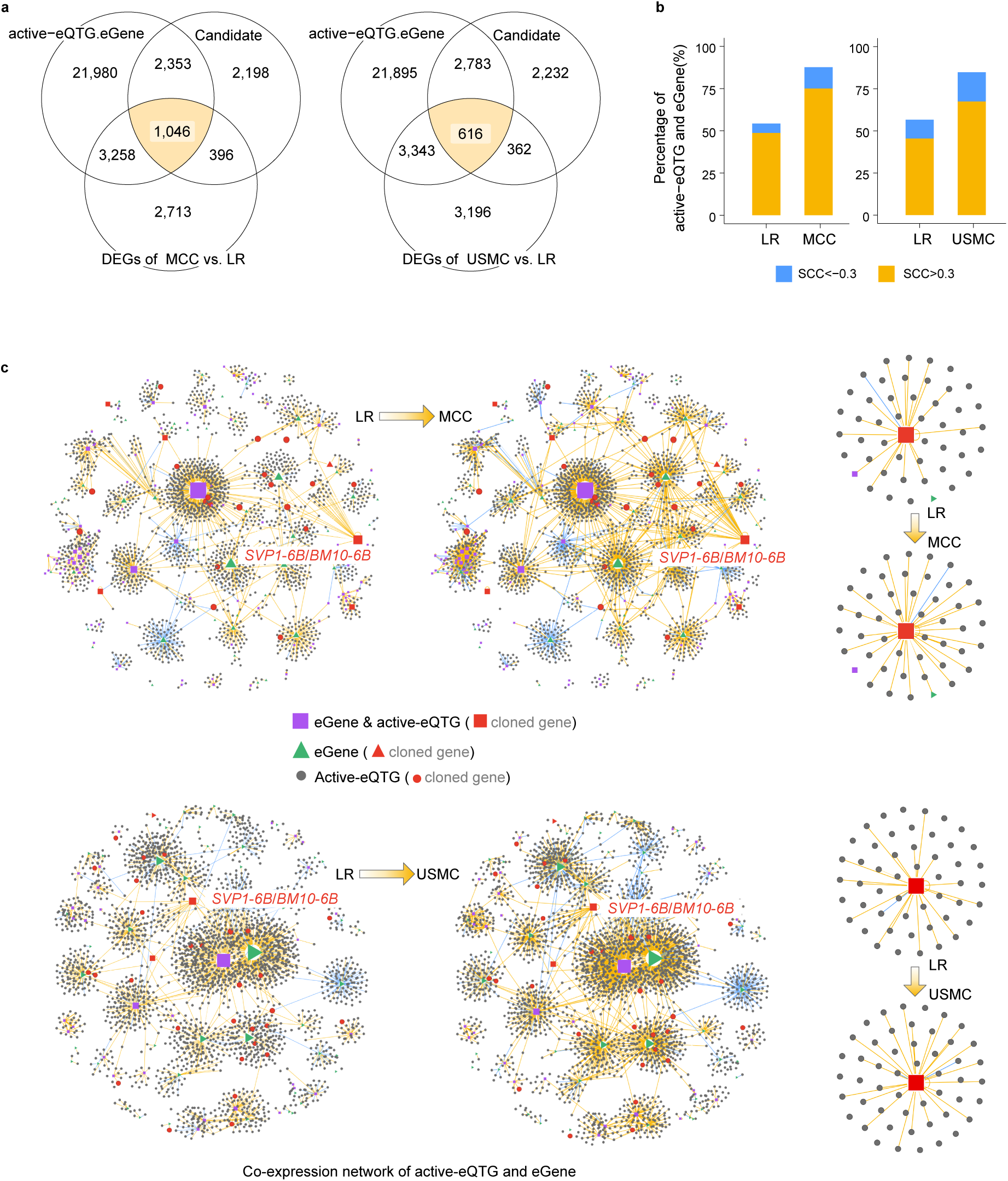
The gene expression networks of different breeding programs. **a** Venn diagram of candidate genes, eGenes regulated by active-eQTGs, and DEGs. **b** The proportion of candidate genes in the orange blocks of the panel (Fig. 7a) and active-eQTGs with Spearman’s rank correlation coefficient |SCC| > 0.3. **c** Network of the genes in the orange blocks from panel (Fig. 7a). The orange lines represent the gene expression correlation (SCC>0.3) between eGenes and active-eQTGs while the blue lines represent the negative correlation (SCC< −0.3). The small regulatory network on the right side displays the co-expression relationship between the cloned genes *SVP1*-*6B*/*BM10*-*6B* and active-eQTGs. MCC is for Modern Chinese Cultivars. USMC is for United States Modern Cultivars. LR is for landraces.

Additionally, among the 297 candidate genes for powdery mildew resistance, 285 candidate genes were regulated by 4,842 active-eQTGs. In the landrace group, MCC group and USMC group, there were 2,900 pairs (59.9%), 3,647 pairs (75.4%) and 3,505 pairs (72.5%) of active-eQTGs and eGenes showing co-expression (|SCC|>0.3), respectively. The MCC and USMC groups had significantly higher proportions of co-expression relationships compared to the landrace group (Fisher’s exact test, *p*-value < 2.2e-16 for both MCC vs. LR and USMC vs. LR) (Supplementary Fig. 16a, b). The powdery mildew resistance gene *Pm4* was predominantly found in the MCC group in this study, resulting in four co-expression regulatory relationships in the MCC network only. The cloned powdery mildew resistance gene *Pm5* had only one co-expression regulatory relationship in the co-expression network (Supplementary Fig. 16c). The simplicity of the co-expression regulatory relationships for these two cloned resistance genes may be attributed to the fact that the traits they control are qualitative rather than complex quantitative traits. These findings also indicate that modern breeding has reshaped co-expression regulatory networks, potentially enhancing the ability to cope with adverse environmental conditions and improving population stability.

## Discussion

Currently, several high-quality wheat reference genomes have been released, however, a pan-genome has not yet ready to be utilized to assist NGS short reads alignment [25]. The frequent hybridization among wheat germplasm, along with introgression from both wild relatives and domesticated progenitors, made it challenging to accurately quantify gene expression levels for wheat RNA-seq samples using a single reference genome alone [25, 27]. In this research, we initially mapped RNA-seq data from 328 wheat samples to the Chinese Spring genome, in order to reveal expression biases across chromosomes. This issue likely stems from the widespread incorporation of large foreign chromosomal segments, such as rye 1RS, *Ae*. *markgrafii*, *T. ponticum* and *T*. *timophevii* in wheat breeding. The single reference genome could not quantify the expression of genes that are not existed in the reference. Thus, in the study, we merged the non-redundant gene models of Chinese Spring with other gene models from other wheat accessions into a pan-gene atlas. This lightweight pan-gene atlas will facilitate a more comprehensive and accurate quantification of gene expression levels before a pan-genome is ready.

Introgression in the wheat genome has been intensively studied in recent years thanks to the decreasing cost of NGS technologies [6, 7, 31, 73], but genome-wide pattern of gene expression for introgression has been rarely studied [24, 25]. Our population-level gene expression analysis revealed a hidden consistency between the expression of introgressed genes and goal of breeding improvement. Genes related to environmental stress or disease resistance are upregulated after introgression while genes of basic cellular process are downregulated or silenced compared to their donor species. Those changes might be achieved through *trans*-eQTL located on other wheat chromosomes. We infer only wheat lines with favorable allele at those *trans*-eQTL locations could allow the optimal expression of introgressed genes. In other words, our calculation suggested the gene expression activity of introgressed allele depends on the genetic background, which is a phenomena frequently observed in distant hybrid breeding [74, 75]. Our results provided clues for designing future breeding plan by selecting hybrid parents with favorable alleles at those *trans*-eQTL.

The integrated pan-gene atlas discovered that 27% transcripts detected in the seedling of a wheat line are not encoded by the Chinese Spring reference genome. In total, more than twenty thousand non-CS transcript were detected for our panel of 328 wheat lines. Although numerous RNA-seq and GWAS have been conducted in wheat, those non-CS genes have been always ignored in previous studies [6, 24, 73]. Using a multi-omic modeling pipeline similar to our previous work [24], we obtained candidate genes of agronomic traits and disease resistant traits, which included 51 non-CS genes for a set of 34 agronomic traits, and 3 non-CS genes for a set of 8 *Bgt* isolates. We used an indexed EMS library to validate if the mutation of a candidate gene has the potential impact on the corresponding trait. Among 72 candidate genes with at least 5 mutant lines, 58 shows significant trait difference between wild type and mutant lines, suggesting our candidate genes predicted by multiple evidence are high value target for further functional investigations.

Compared to the eQTL analysis performed in our previous work [24], the presented work used WGS derived SNP data. Thus, the regulatory elements located far from gene region were fully covered in this work. Although intergenic SNP was sometimes removed in genomic studies [73], they do play a role in gene regulation. In order to reveal their role in gene regulation, we merged significant SNPs from gene and non-gene regions into eQTLs and found that non-gene region eQTLs play an important role in gene regulation and the prediction of candidate genes for agronomic traits. The genic region of 2%-5% candidate genes did not show any GWAS signal. The association of expression for those genes with agronomic traits are mediated through an eQTL from inter-genic region. So, the SNP in inter-genic region is important for us to understand the genetic and molecular mechanism of agronomic traits. Marker assisted breeding could utilize those inter-genic eQTLs.

Modern breeding not only involves the selection of target genes but also changed transcriptome profiles at the population level. Approximately 10% of genes exhibit differential expression between different sub-populations. Notably, differences exist in the direction of changes between the cultivars from the China and that from the United States, which is likely attributed to varying breeding objectives and environmental adaptability. Despite some shared gene regulatory changes between those two countries, such as key genes involved in environmental stress responses and photoperiod regulation (e.g., *VRN1*, *VRN2*, *WRKY45*), the differing breeding objectives may led to distinct gene expression patterns. For example, breeders in the United States emphasize genes that improve nitrogen use efficiency and minimize fertilizer usage (e.g., *TaGS1a-6B*) [70], while breeders in China favor genes of disease resistance and yield (e.g., *Lr34*) [64]. These differing directions of gene expression changes reflect the difference of breeding objectives between countries, as well as the varying selection strategies adopted to address the unique challenges faced in their agricultural production. Furthermore, we found that DEGs not only exhibit greater genetic differentiation compared to non-DEGs, but are also subjected to stronger selective pressure, a pattern that extends to the regulatory sites controlling these genes. Thus, modern breeding has significantly altered the gene expression regulatory network in wheat, impacting the expression patterns of thousands of genes. These expression differences not only reflect the breeding goals of different countries but also reveal unique genetic regulatory patterns shaped by selection pressures in different environmental conditions. Future studies can further investigate the functions of these DEGs in specific agronomic traits.

Changes in gene expression during breeding process are not only limited to individual genes but also involve more complex shifts in gene co-expression and regulatory networks. Our study found that the number of co-expression pairs involving identified active-eQTGs and eGenes is significantly lower in landraces compared to cultivars. This suggests that modern breeding has reshaped the regulatory networks of cultivars, potentially by selecting genes related to environmental adaptation, thereby altering the overall gene co-expression network. These network changes may contribute to the enhanced stability of agronomic traits and better adaptation to adverse environmental conditions in cultivars. Therefore, future breeding strategies should not only focus on the selection of specific genes but also on optimizing and utilizing the overall regulatory networks.

## Materials and methods

### Wheat germplasm and phenotypes

We selected 328 accessions from previously published whole-genome resequencing data of 355 common wheat (*T. aestivum*) [6], including 92 modern Chinese cultivars (MCC), 64 modern United States cultivars (USMC), and 172 landraces (LR) from 13 countries worldwide, representing a wide range of genetic diversity (Supplementary Table 1).

We evaluated 34 field agronomic traits and seedling-stage resistance phenotypes of 8 *Blumeria graminis* f. *sp*. *Tritici* (*Bgt*) isolates across the 328 wheat accessions. The planting and phenotypic measurements of field agronomic traits were carried out simultaneously with those reported in previous studies. All 328 common wheat accessions were grown during three consecutive winter growing seasons (2013 to 2016) in Zhao County, Shijiazhuang City, Hebei Province, China (38° 05′N, 114°52′E). The phenotypic data for 20 of these traits have already been published. These include four grain-related traits: Grain length (GL), Grain roundness (GRO), Grain width (GW), and Grain number (GN); seven yield-related traits: Thousand kernel weight (TKW), Harvest index (HI), Yield per plant (YPP), Spikelet number (SPN), Awn length (AL), Sterile spikelet number (SSN), and Biomass per plant (BPP); and nine growth and development traits: Anthesis days (AD), Heading days (HD), Flag leaf length (FLL), Flag leaf width (FLW), Plant height (PH), Peduncle length (PL), Stem diameter (SD), Tiller number at jointing stage (TNJS), and Tiller number at seedling stage (TNSS) [6]. 14 new field agronomic trait phenotypes are included in this study, consisting of seven grain-related traits: Grain filling days (GFD), Grain area (GA), Grain diameter (GD), Grain perimeter (GP), Grain red (GR), Grain green (GG), and Grain blue (GB); five yield-related traits: Yield per head per plant (YHPP), Biomass per plant at jointing stage (BPPJS), Biomass per tiller at jointing stage (BPTJS), Biomass per plant at seedling stage (BPPSS), and Biomass per tiller at seedling stage (BPTSS); and two growth and development traits: Lifecycle (LC) and Days from heading to anthesis (HAD). The Best Linear Unbiased Estimates (BLUEs) were derived using a model with fixed genotype effects and random effects for each year.

We isolated and purified eight *Bgt* isolates from fields across different provinces in China, naming them B040A1, B056A1, B080A1, B094A1, B099A1, B114A2, B132A2 and B138A1 (Supplementary Table 9). We planted the 328 wheat accessions in rectangular trays with three replicates, providing 14 hours of light at 22 °C and 10 hours of darkness at 18 °C. We inoculated the plants with the *Bgt* isolates at the one-leaf stage for each of the 8 isolates. Infection types were evaluated 10 days after inoculation using a scale ranging from 0 to 4, where 0 indicates necrotic flecks, and 1 to 4 represent highly resistant, moderately resistant, moderately susceptible, and highly susceptible responses, respectively [52].

### RNA-seq sequencing and data preprocessing

A total of 328 wheat lines were grown in trays, with each line having three biological replicates, maintained under a 14-hour light and 10-hour dark cycle at 22 °C and 18 °C, respectively. Leaf tissues were sampled at the two-week seedling stage, and the leaves from the three biological replicates were averaged for RNA extraction using the FastPure Universal Plant Total RNA Isolation Kit. The extracted RNA was then used to construct libraries via the BGI Optimal Series Dual Module mRNA Library Construction Kit (LR00R96) and sequenced on the DNBSEQ-T7 platform, generating 2 × 150 bp reads. In parallel, 7 lines of *T. ponticum* were also grown under identical conditions for RNA sequencing. Additionally, we obtained published RNA data from previous studies, which included leaf samples from 10 rye samples, 2 Renan samples, and 3 *T. timophevii* samples (Supplementary Table 6), all at the 2-3 week growth stage. The software Salmon (v1.8.0) [76] program that utilizes pseudoalignment techniques on RNA-seq reads to reference gene models was used to quantify the transcript abundance. program that utilizes pseudoalignment techniques on RNA-seq reads to reference gene models was used to quantify the transcript abundance.

### Identification of large introgression events using WGS genotyping data

The genotypes of 328 wheat accessions were extracted from the original high-density genotype VCF file from a previous study of 355 wheat accessions using the Chinese Spring reference genome (IWGSC RefSeq v1.1). After filtering out variants with a missing rate > 25% and heterozygosity > 30%, variants with MAF > 0.05 were retained for this study, resulting in a total of 26,788,626 variants, including 24,744,215 SNPs and 2,044,411 InDels.

To detect non-CS introgression fragments, a whole-genome genotype heatmap was generated using ComplexHeatmap [77, 78], with a window size of 1 SNP/MB, revealing missing segments >20 Mb. To further quantify these deletions, we calculated the missing rate across the genome using a 2 Mb sliding window with a 1 Mb step size, then compared the missing rates in deleted versus non-deleted regions, and tested for significance using the Wilcoxon test.

### Reference genome model of large introgression segments

In this study, the four large introgressed segments have been previously reported, so the genomes of these donors were used as reference models. We used the genomes of *Secale cereale L*., the French cultivar Renan (Instead of *Ae. markgrafii*, because the terminal end of Renan 2D has been reported to possibly originate from the introgression of *Ae. markgrafii* without a published genome), *Thinopyrum elongatum* (Instead of *T. ponticum*, because the genome of *T. ponticum* has not been published, and *Th*. *elongatum* is closely related to *T*. *ponticum*), and *Triticum timopheevii*, These genomes were merged with the CS RefSeq v1.1 reference genome. Then, based on the merged gene models, gene expression levels of 328 wheat samples were calculated using Salmon (v1.8.0) [76]. The number of expressed genes with expression (TPM > 0.5) was then counted in 10 Mb windows with a 2.5 Mb step along each chromosome.

### The construction of pan-gene atlas

Beyond greater than 20 Mb introgressed fragments, which can be easily identified through genotype missing rate calculations, smaller chromosome segments cannot be accurately evaluated using the Chinese Spring gene model as a reference. To address this, we employed gene models from 24 published *Triticeae* genomes, including hexaploid species such as Chinese Spring (IWGSC RefSeq v1.1, 2n=6x=42, AABBDD), Kenong9204 (KN9204) [77], Aikang58 (AK58) [78], Fielder [79], and 10 wheat genomes (ArinaLrFor, Jagger, Julius, LongReach Lancer, CDC Landmark, Mace, Norin 61, CDC Stanley, SY Mattis, and Spelt) [29, 82]. We also used tetraploid genomes, including *T. turgidum* ssp. *dicoccoides* (wild emmer, 2n=4x=28, AABB) and *Triticum turgidum* L. ssp. *durum* (durum, 2n=4x=28, AABB) [83], along with diploid genomes such as *Triticum urartu* (2n=2x=14, AA) [84], *Triticum monococcum* (2n=2x=14, AA) [85], *Aegilops speltoides* (2n=2x=14, SS) [86], and *Aegilops tauschii* (2n=2x=14, DD) [87]. And 4 gene models from large introgressed segments such as *Secale cereale L*. (rye, chr1R: 0-280 Mb), the French cultivar Renan (chr2D: 570-635 Mb) (Instead of *Ae. markgrafii*), *Thinopyrum elongatum* (chr3E: 500-676 Mb) (Instead of *T. ponticum*), and *Triticum timopheevii* (2n=4x=28, A^t^A^t^GG, chr5G: 435-485 Mb) to construct a pan-gene atlas.

The construction of the pan-gene atlas consists of three parts. The first part includes all high-confidence (HC) genes from Chinese Spring (CS). The second part consists of all HC genes from four large introgressed segments: Rye (chr1R: 0-280 Mb), Renan (chr2D: 570-635 Mb), *Th. elongatum* (chr3E: 500-676 Mb), and *T. timopheevii* (chr5G: 435-485 Mb). The third part contains non-redundant HC genes from 13 hexaploid genomes, 2 tetraploid genomes, and 4 diploid genomes. We first utilized OrthoFinder (v2.5.4) [88, 89] to identify the orthogroups among all genes from Chinese Spring and the 19 genomes. Orthogroups containing genes from Chinese Spring were represented by the Chinese Spring genes, thus these Orthogroups were excluded. For orthogroups without Chinese Spring genes, the longest transcript was retained as the representative gene. The genes from these three parts were merged into a pan-gene atlas. We used the pan-gene atlas as a reference and employed Salmon (v1.8.0) [76] to assess the transcript abundance of 328 wheat samples through pseudoalignment, which was used for all subsequent analyses. The construction of the pan-gene atlas consists of three parts. The first part includes all high-confiden.

### Quantification of gene expression for introgressed segments

We used OrthoFinder (v2.5.4) [88, 89] to identify homologous high-confidence (HC) genes between rye (chr1R: 0–280 Mb) and Chinese Spring (CS). Homologous genes between rye and wheat are termed conserved genes, while those unique to rye are called specific genes. In order to analyze the relationship between the similarity of rye 1RS conserved genes and their expression ratio in wheat, we filtered the 1RS genes based on alignments to wheat genes with a length greater than 80% and identity over 70%.

We performed differential expression analysis of chromosome 1RS genes between the 1RS.1AL, 1RS.1BL introgression lines and the 10 rye accessions. First, we transformed the gene expression levels using log2(TPM+1), followed by normalization of the expression matrix using the normalize.quantiles.robust method. Next, we analyzed differentially expressed genes between the 1RS.1AL introgression line and rye chromosome 1RS, as well as between the 1RS.1BL introgression line and rye 1RS, using two methods: DESeq2 and the Wilcoxon rank-sum test [90]. In DESeq2, the criteria for defining differentially expressed genes were *P*adj < 0.05 and abs(log2FoldChange) > 1. For the wilcoxon rank-sum test, log2FoldChange was calculated as the fold change in average gene expression between group 1 and group 2, with differentially expressed genes defined by thresholds of *P*adj < 0.05 and abs(log2FoldChange) > 0.5. These same criteria were applied for the analysis of other introgressed genes.

We utilized InterProScan (v. 5.66-98.0) [91] to annotate the genes of rye, diploid *Th. elongatum*, and *T. timopheevii*. The GO enrichment analysis for differentially expressed genes was performed using TBtools [92].

### Detection of eQTL

We performed the association analysis between SNPs and gene expression PEER residuals using the R package MatrixEQTL with useModel = modelLINEAR [93]. The first five principal components (PCs) of the SNP matrix, representing population structure, were included as covariates. SNPs with a false discovery rate (FDR) threshold < 1e-10 were considered significant signals associated with gene expression.

SNPs associated with gene expression are divided into three types. First, SNPs were annotated using SnpEff (v.5.0e) [94]. and classified as intergenic SNPs or genic SNPs. Genic SNPs were further classified based on whether the genes they are located in are expressed. If a gene is expressed in more than 5% of the population samples (TPM > 0.5), it is considered as an active gene, and the corresponding SNPs are labeled active SNPs. Genes not expressed at the population level are defined as inactive genes, and their SNPs are categorized as inactive SNPs. Subsequently, we merged the significant SNPs of the three types using strict criteria. Intergenic SNPs were merged based on the criteria of continuous SNPs within < 100 kb, a minimum of three SNPs, and LD > 0.4, retaining the most significant SNP as the intergenic-eQTL. For inactive SNPs, the merging strategy involved first combining them based on intergenic regions, retaining the most significant SNPs, followed by merging based on LD > 0.4, with the most significant SNP retained as the inactive-eQTG; active SNPs were merged by first retaining high-effect SNPs within the gene. If there were more than one SNP, the SNP with the strongest signal was retained as the active-eQTG. Second, the merging process for active SNPs is as follows: we first compute the SCC correlation of expression levels between all active genes and their regulatory eGenes across genotypes. For active genes with high-effect SNPs, the most significant SNP is retained as the active-eQTG. For non-high-effect SNPs, we consider active genes with |SCC| > 0.3 to be correlated with their eGenes, retaining the most significant SNP on these genes as the active-eQTG. Finally, we retain both high-effect active-eQTGs and those with |SCC| > 0.3. Intergenic-eQTLs, inactive-eQTGs, and active-eQTGs are collectively referred to as eQTLs. An eQTL and its eGene located on the same chromosome are classified as *cis*-eQTLs, while those on different chromosomes are classified as *trans*-eQTLs. This classification applies only to CS eGenes.

### GWAS

A GWAS was performed using a linear mixed model that addressed both population structure and kinship for all 34 field agronomic traits and 8 seedling-stage *Bgt* isolate phenotypes, employing the ‘--mlma’ parameter in GCTA (v1.94.1) [95]. The first five principal components were used to control for population structure. A kinship matrix that represents pedigree relationships was generated using a subset of independent SNPs in GCTA. If two consecutive significant SNPs were located less than 2 Mb apart, they were classified as a single QTL, with the most significant SNP retained as the representative signal for the QTL. Genes that contained significant SNP variations within the QTL interval were designated as candidate genes for the QTL region.

### TWAS

As described in a previous study [45], transcriptome-wide association studies (TWAS) provide a framework for identifying significant *cis* genetic variants correlations between gene expression and phenotype. For this research, the Fusion program was utilized to carry out the TWAS (http://gusevlab.org/projects/fusion/). The program requires the computation of gene expression weights, reflecting the pre-modelled relationships between SNPs and gene expression levels, which are then integrated with GWAS to estimate the associations between genes and phenotypic traits.

We selected the SNPs located within 2 Mb upstream and downstream of the Chinese Spring (CS) genes and their corresponding expression levels as phenotypic traits to estimate the expression weights of the CS genes at the population level. For non-CS genes, we identified the physical positions based on the most significant eQTL signals in the Chinese Spring genome, using the SNPs and expression levels within a 2 Mb range to compute their expression weights. Heritability calculations were carried out using GCTA (v1.94.1) [95], and expression weights were derived from models such as top1, blup, lasso, and enet. Subsequently, we extracted the SNP data from the GWAS results, including A1 (first allele) and A2 (second allele), and computed the Z-scores with the formula Z-scores = beta/se. The FUSION.test.R script from the Fusion program was employed to assess gene signals associated with phenotypic traits across each chromosome. Genes with TWAS *p*-value < 1×10^−3^ were considered candidate genes.

### SMR

We performed a summary data-based Mendelian randomization analysis (SMR) to investigate the association between gene expression and trait variation, utilizing summary-level data from our eQTL mapping study and GWAS results for 34 field agronomic traits and 8 seedling-stage *Bgt* isolates phenotypes, employing GCTA (v1.94.1) [95]. For the physical location information of non-CS genes, we approximated the positions of the strongest eQTL signals from CS as the physical locations of the non-CS genes. The analysis of summary-level statistic of these two GWAS datasets was conducted using the SMR commands ‘-cis-wind 10000’ for cis-eQTL and ‘--trans-wind 5000’ for trans-eQTL.Genes with SMR *p*-value < 1×10^−4^ were designated as candidate genes.

### Validation of candidate genes using the EMS mutant library

Genes in KN9204 with a sequence similarity greater than 98% and coverage greater than 90% to the candidate genes were identified as homologous genes. Homologous genes with at least five non-synonymous mutants in the indexed KN9204 EMS library [53] were selected for further analysis. A two-tailed Student’s *t*-test was conducted between the mutants (n > 5) and wild-type samples (n = 15). Considering the random nature of mutations in the EMS library, we introduced a control by randomly selecting an equal number of genes without mutations in the candidate genes to validate the accuracy of the tests. We evaluated the effects of candidate gene mutations on seven agronomic traits, including Spikelet number (SPN), Grain number (GN), Grain area (GA), Grain perimeter (GP), Grain length (GL), Grain width (GW), Thousand kernel weight (TKW). Given the high Pearson correlation coefficient among 34 field agronomic traits (Supplementary Fig. 9), mutations in candidate genes affecting one trait often influence multiple traits. Thus, candidate genes associated with these agronomic traits were tested across all seven selected traits.

### Detection of DEGs between cultivars and landraces

We performed differential expression gene analysis on three populations: MCC, USMC, and LR, employing the Wilcoxon rank-sum test. The normalization of gene expression levels follows the same method as described above. The thresholds for identifying DEGs were *P*adj < 0.05 and |log2FoldChange| > 0.5, where log2FoldChange represents the ratio of the mean gene expression levels between population 1 and population 2. DEGs were categorized into four types across eight models, using the landraces gene as a reference. The first category (M1 and M2) includes genes with inconsistent regulation between MCC and USMC relative to landraces; the second category (M3 and M4) includes genes with consistent regulation. The third category (M5 and M6) consists of differential genes between MCC and landraces, with no significant differences for USMC and landraces. The fourth category (M7 and M8) consists of differential genes between USMC and landraces, with no significant differences for MCC and landraces. Finally, GO enrichment analysis was conducted for the 8 models of DEGs using TBtools [92].

### Dimensionality reduction of population expression

Following the pairwise differential expression analysis of the populations MCC, USMC, and landraces (LR) we identified unique differentially expressed genes for use in the dimensionality reduction analysis. The standardization method for the expression levels of the DEGs was consistent with the previously described approach. Subsequently, we employed the umap function from the R package uwot to perform Uniform Manifold Approximation and Projection (UMAP) for dimensionality reduction analysis [96].

### Population genetics analysis

Given the large LD distance and dense SNP coverage, we filtered SNPs based on the criteria from the published work [6], retaining 5,749,696 SNPs/InDels for PCA analysis using PLINK (v1.9) [97]. The genetic differentiation (*F*_ST_) between subpopulations (MCC vs. LR, USMC vs. LR, USMC vs. MCC) were calculated using a 20-kb sliding window and a step size of 10 kb with VCFtools (v0.1.16) [98]. To assess the genetic differentiation between DEGs, non-DEGs, and regulatory regions in modern breeding, we compared the *F*_ST_ values for the respective regions. To analyze the relationships between the signals of DEGs and the most significant eQTLs signals with *F*_ST_, we paired the signals of DEGs and the most significant eQTLs signals with their corresponding *F*_ST_, sorting them by *F*_ST_. A custom script was used to split the data into ten bins, calculating the mean and standard error of the signals of DEGs and top eQTL signals as *F*_ST_ changes.

### Detection of selective sweeps between landraces and cultivars

We utilized the Python-based composite likelihood approach (XP-CLR) [99] to identify selective sweeps that occurred during the improvement of modern breeding (https://github.com/hardingnj/xpclr). For this analysis, landraces were considered as the reference group, with MCC and USMC acting as the query groups. We scanned for selective sweeps with a step size of 10 kb and a 20 kb sliding window across each chromosome (--size 20,000;--step 10,000). We obtained XP-CLR scores for the regions associated with DEGs to evaluate whether they were subjected to selection pressures during modern breeding processes. Moreover, we also extracted XP-CLR scores for the regions of the most significant eQTLs of DEGs to ascertain whether these regulatory regions were influenced by breeding selection. We filtered out the sites with XP-CLR scores equal to zero, sorted the remaining data by XP-CLR scores, and divided the results into 20 equal bins. Using a custom script, we calculated the mean and standard error of the signals for both differential expression and eQTLs in each bin.

### Construction of gene co-expression regulatory networks

Active-eQTGs regulate the expression of eGenes, and both active-eQTGs and eGenes are expressed in the population. Therefore, we utilized this characteristic to construct co-expression regulatory networks. We first obtained the relationships between the selected eGenes and their corresponding active-eQTGs, then calculated the Spearman Correlation Coefficients (SCC) of eGenes and active-eQTGs within sub-populations. |SCC| > 0.3 were considered as co-expression pairs. We then counted the number of co-expression relationships in each sub-population. We visualized the regulatory relationships between these active-eQTGs and eGenes using Gephi (https://gephi.org/), with orange edges representing positive correlations, blue edges representing negative correlations, and no edges displayed for regulatory relationships without co-expression.

## Authors’ contributions

F.H. and H.-Q.L. designed and supervised the research, and edited the manuscript. Z.Z. leads the data analysis and drafted the manuscript with the input from M.S. M.Y. contributed to the overall strategy of data analysis and interpretation of the results. C.Z. provided the 8 *Bgt* isolates phenotypes. Y.Y. and X.Z. conducted the qRT-PCR experiments. J.X. participated in part of the bioinformatics analysis. Y.S., L.W., and X.Z. contributed to the interpretation of gene function. Q.Z. provided materials for *T. ponticum*. S.Z. and H.-Q.L. supplied field agronomic traits data for our RNA-seq panel. H.W., X.L. and N.J. provided trait for EMS mutant lines. J.W., Y.Z. helped editing the manuscript. All the authors reviewed and approved the manuscript.

## Funding and Acknowledgement

This work was supported by the National Key Research and Development Program of China (2023YFF1000100 and 2024YFE0115100), the Biological Breeding-National Science and Technology Major Project (2023ZD04073), the National Natural Science Foundation of China (32472130 and U24A20391) and the Yazhouwan National Laboratory project (2310JM01).

## Availability of data and materials

The raw sequence data reported in this paper have been deposited in the Genome Sequence Archive [100] of the National Genomics Data Center [101], China National Center for Bioinformation/Beijing Institute of Genomics, Chinese Academy of Sciences (GSA: CRA022106 and CRA022107). The results, including RNA-seq quantification, the pan-gene atlas, orthologous genes and eQTLs, have been uploaded to the Figshare database (https://figshare.com/s/0e907a409ac93a4b4fea).

---

**Supplementary Fig. 1.**
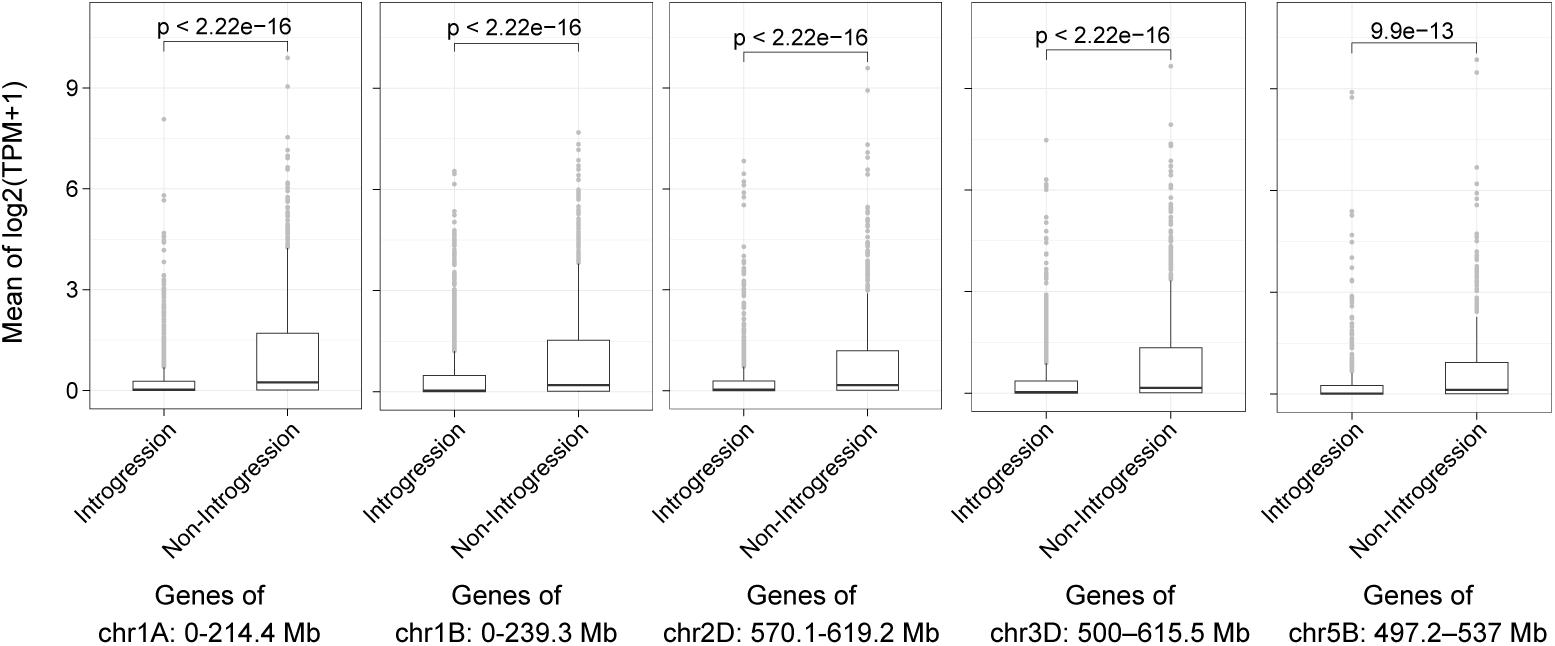
Comparison of the gene expression levels between wheat lines with and without the introgression using Chinese Spring reference genome.

**Supplementary Fig. 2.**
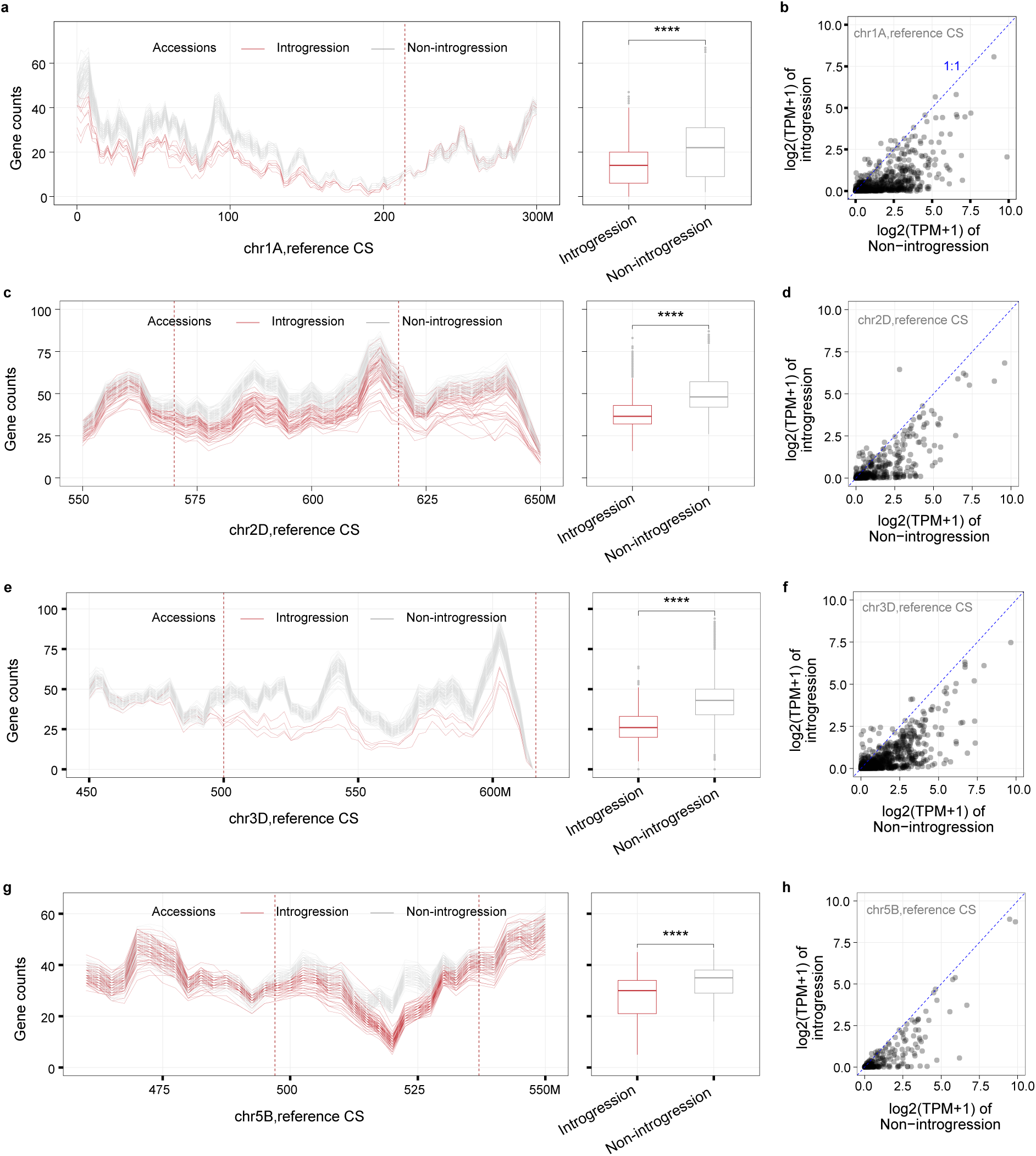
Assessing reference bias in alien introgression of wheat. **a,c,e,g** The left panel shows the number of expressed genes (TPM > 0.5) of wheat lines with and without introgression along chromosome 1AS (0-300 Mb), chromosome 2D (550-650 Mb), chromosome 3D (450 Mb-chromosome end), and chromosome 5B (450-550Mb). A sliding window of 10 Mb with a step size of 2.5 Mb were used. The right panel presents boxplots comparing the number of expressed genes between wheat lines with and without introgression (Wilcoxon rank sum test, \*\**p* < 0.01; \**p* < 0.05). **b,d,f,h** Average gene expression between introgression lines and non-introgression lines in chromosome 1A (0-214.4 Mb), 2D (570.1-619.2 Mb), 3D (500-615.5 Mb) and 5B (497.2–537 Mb). The Chinese Spring genome was used as the only reference for RNA-seq reads mapping here.

**Supplementary Fig. 3.**
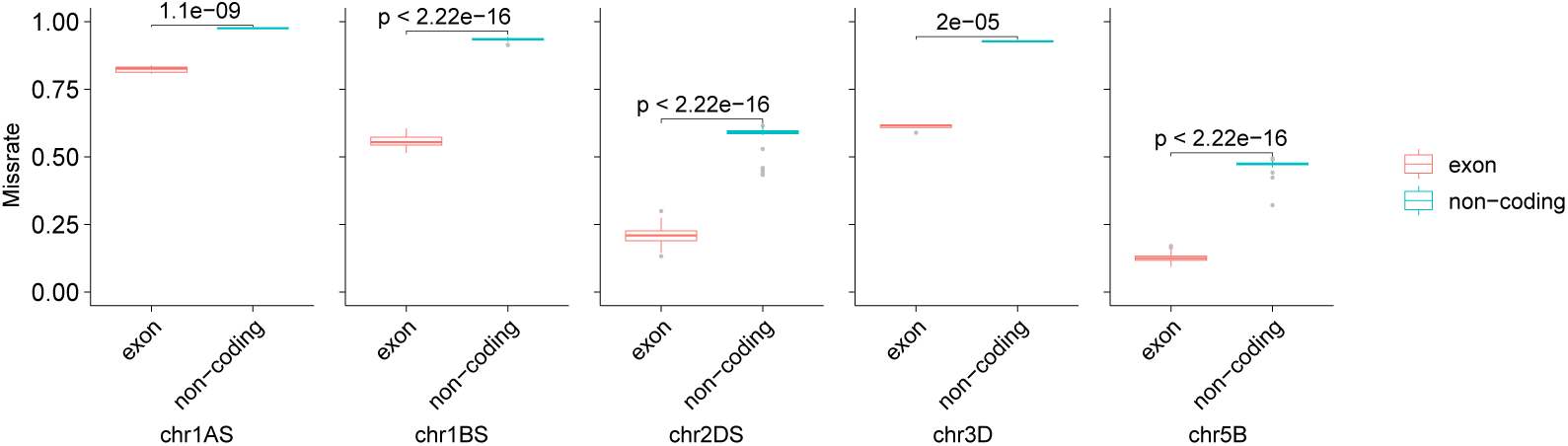
Genotype missing rates in large introgressed segments within exome and non-coding regions of translocation lines calculated using the orginal VCF file of Niu et al, where Chinese Spring was used as the only reference genome.

**Supplementary Fig. 4.**
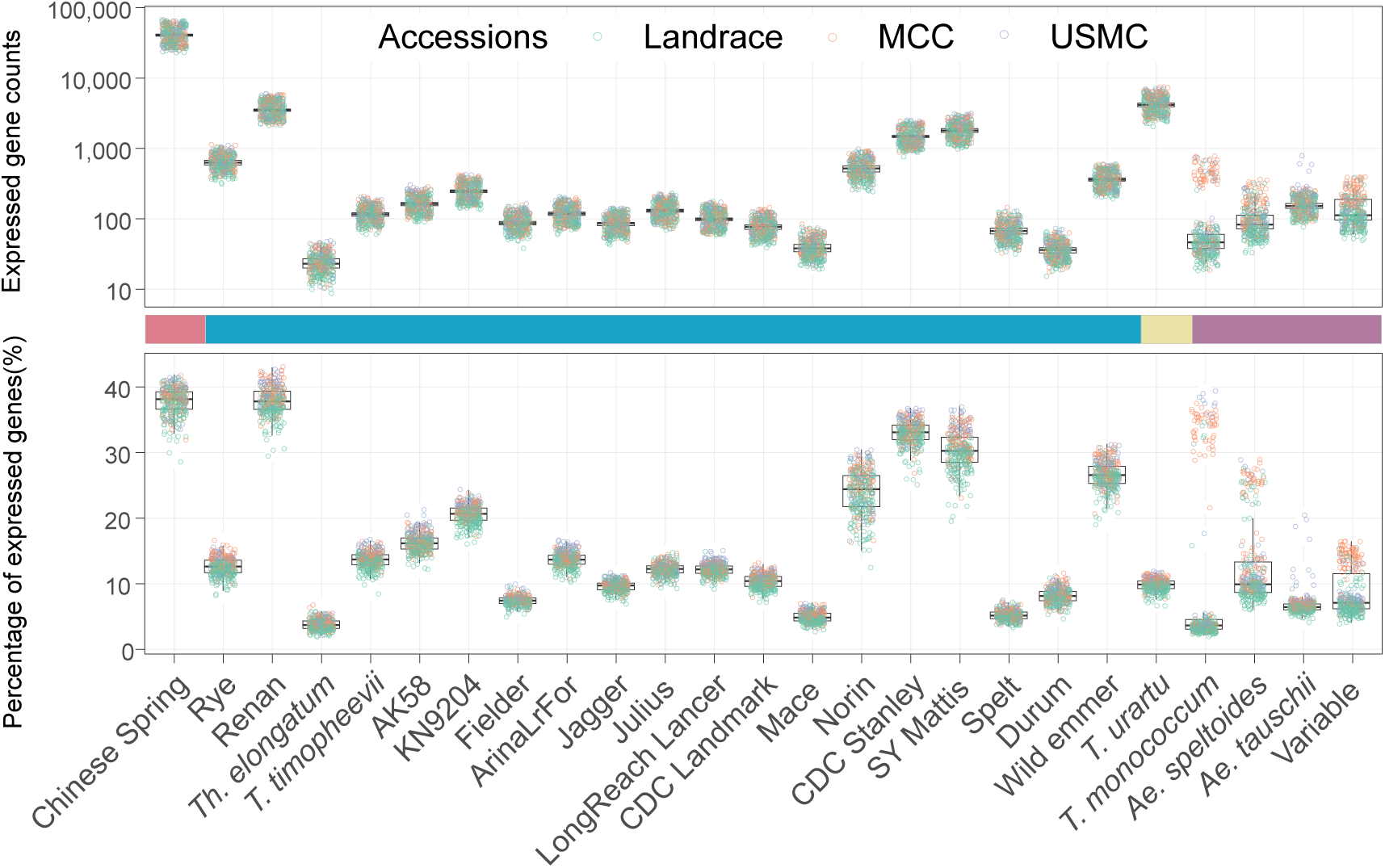
The number and proportion of expressed genes in each genome of the pan-gene atlas for each wheat line of our RNA-seq panel.

**Supplementary Fig. 5.**
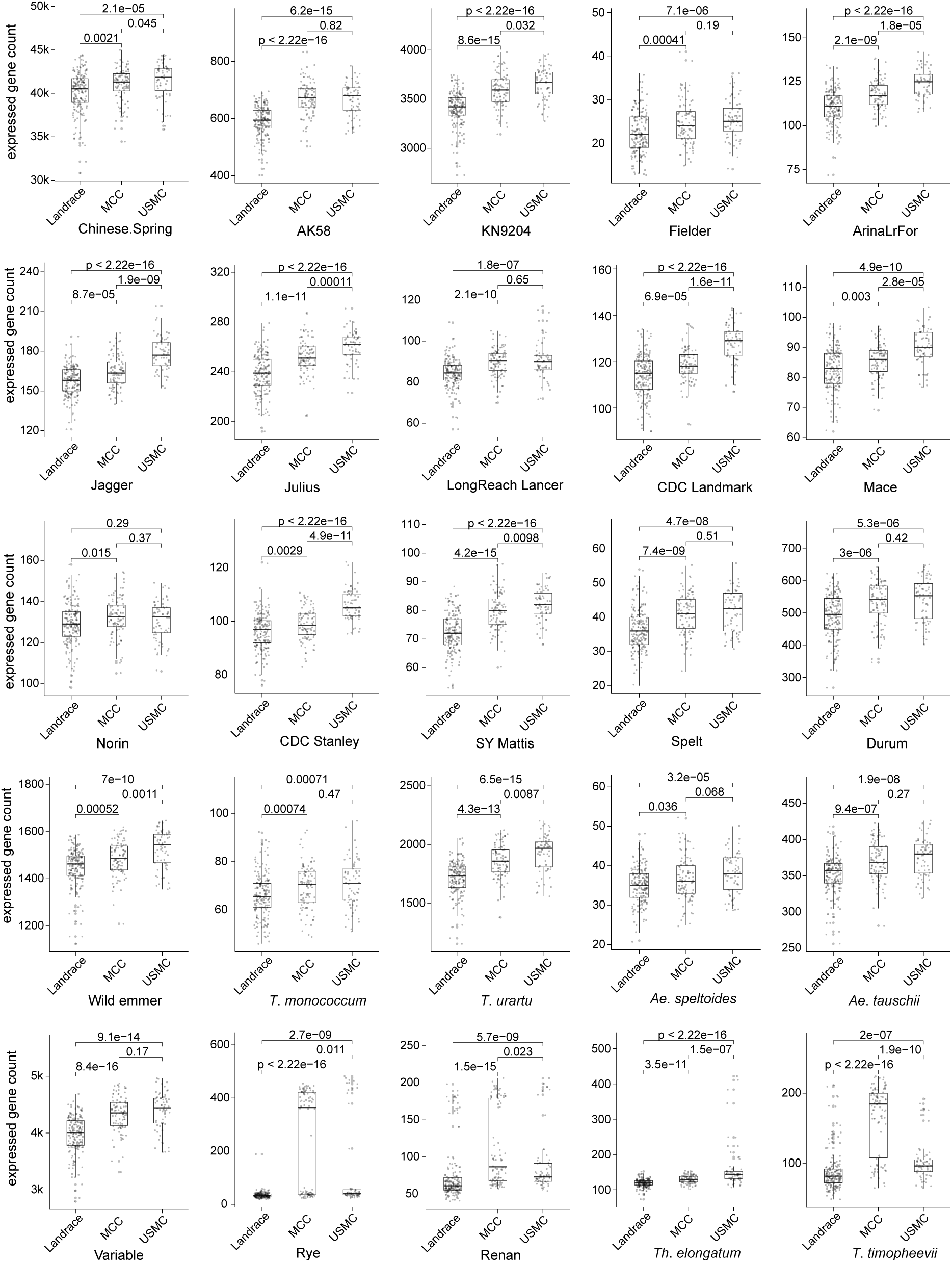
The number and proportion of expressed genes in each genome of the pan-gene atlas for each wheat line grouped by three sub-populations of our RNA-seq panel, including MCC(Modern Chinese Cultivar), USMC(United States Modern Cultivar), and Landrace.

**Supplementary Fig. 6.**
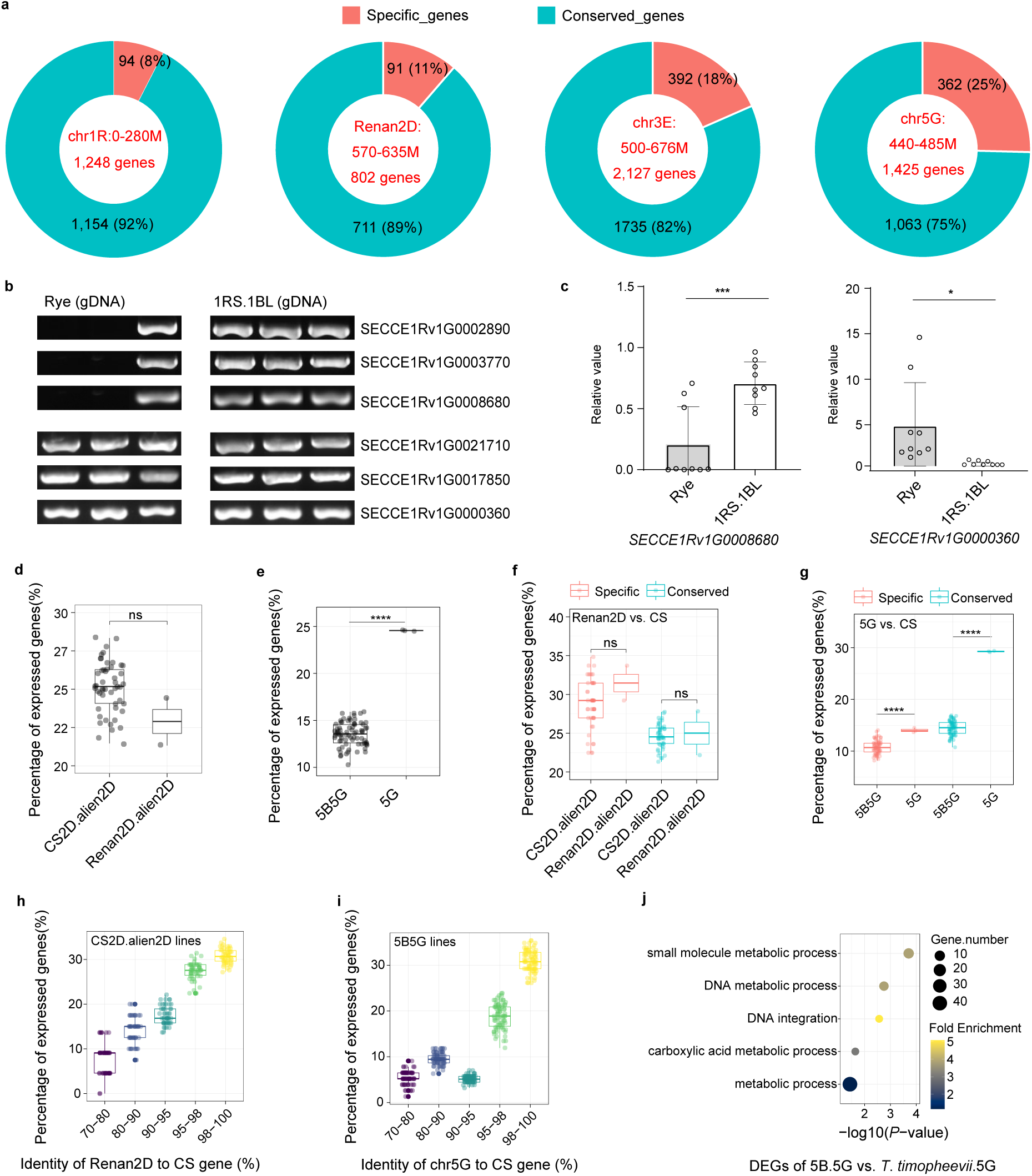
Expression pattern of distantly introgressed genes in wheat and their donor species. **a** The number and proportion of conserved and species-specific genes between distantly introgressed segments and wheat. **b** Amplification of six rye genes in gDNA from three different rye lines and three different 1RS.1BL introgression lines. **c** Transcript-level validation of differentially expressed genes between the 1RS.1BL translocation lines and rye lines using qRT-PCR, with *Tatublin* as the internal control. \**p* < 0.05, \*\**p* < 0.01, \*\*\**p* < 0.001, \*\*\*\**p* < 0.0001 (two-tailed Student’s *t*-test). d The percentage of expressed introgressed genes of *Ae*. *markgrafii* in translocation lines Renan2D, and *Ae*. *markgrafii* translocation lines in this study. **e** The percentage of expressed genes in the chromosome 5G (440-485 Mb) region in *T*. *timophevii* translocation lines compared to *T*. *timophevii* lines. **f,g** Comparison of the proportion of expressed genes between conserved and specific introgressed genes in the translocation lines and donors. **h,i** The number of expressed genes in the introgression line grouped by the degree of sequence conservation between wheat and the wild relatives. **j** GeneOntology enrichment analysis of differentially expressed introgressed genes between the *T*. *timophevii* translocation lines and *T*. *timophevii* lines.

**Supplementary Fig. 7.**
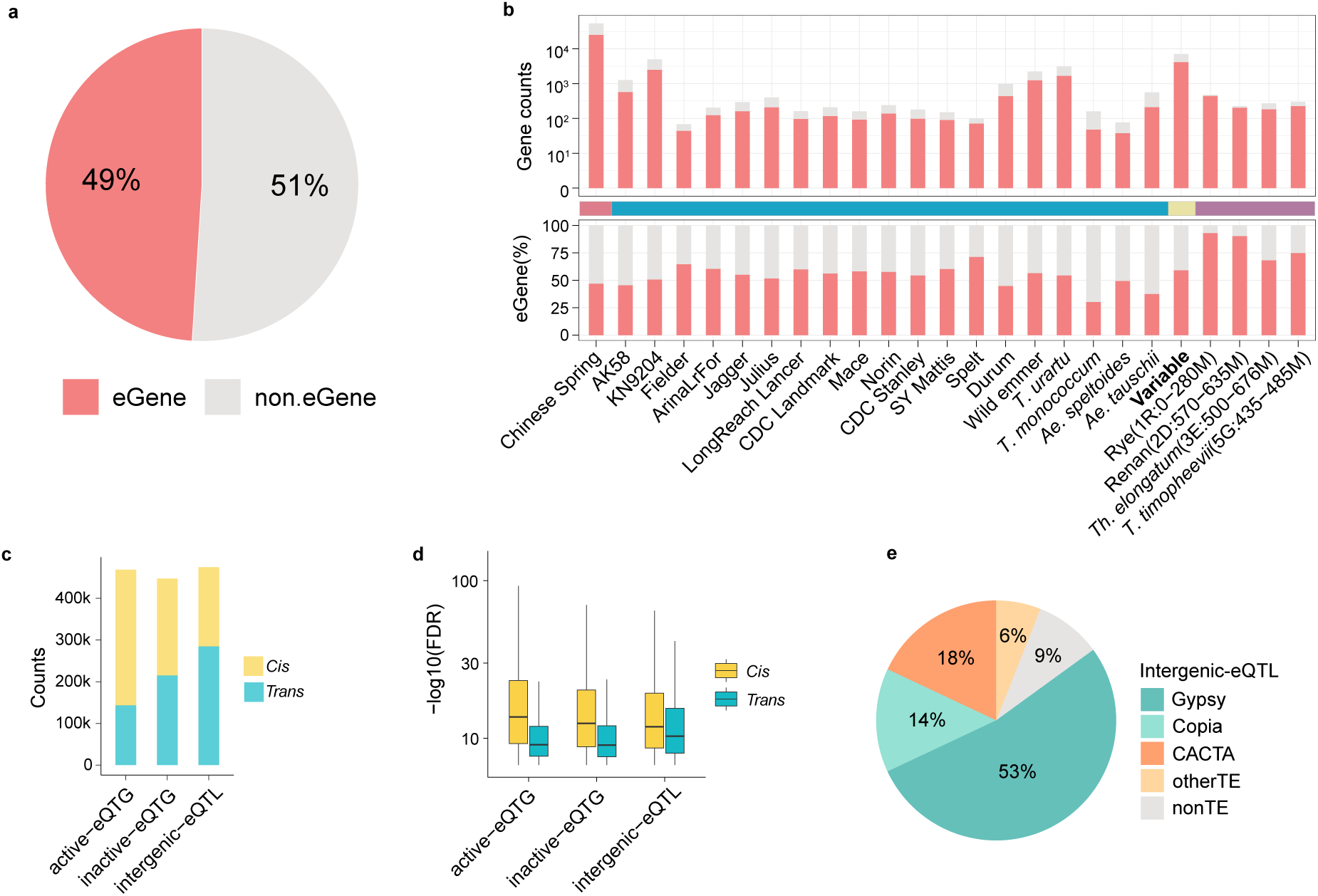
The eQTL feature of Pan-gene atlas. **a** The proportion of eGenes in the Pan-gene atlas. **b** The number and proportion of eGenes in 24 genomes of the Pan-gene atlas. **c** The number of *cis*- and *trans*-eQTLs. **d** The strength of association signals for *cis-* and *trans*-eQTLs. **e** The classification and proportion of intergenic-eQTLs in the transposon region.

**Supplementary Fig. 8.**
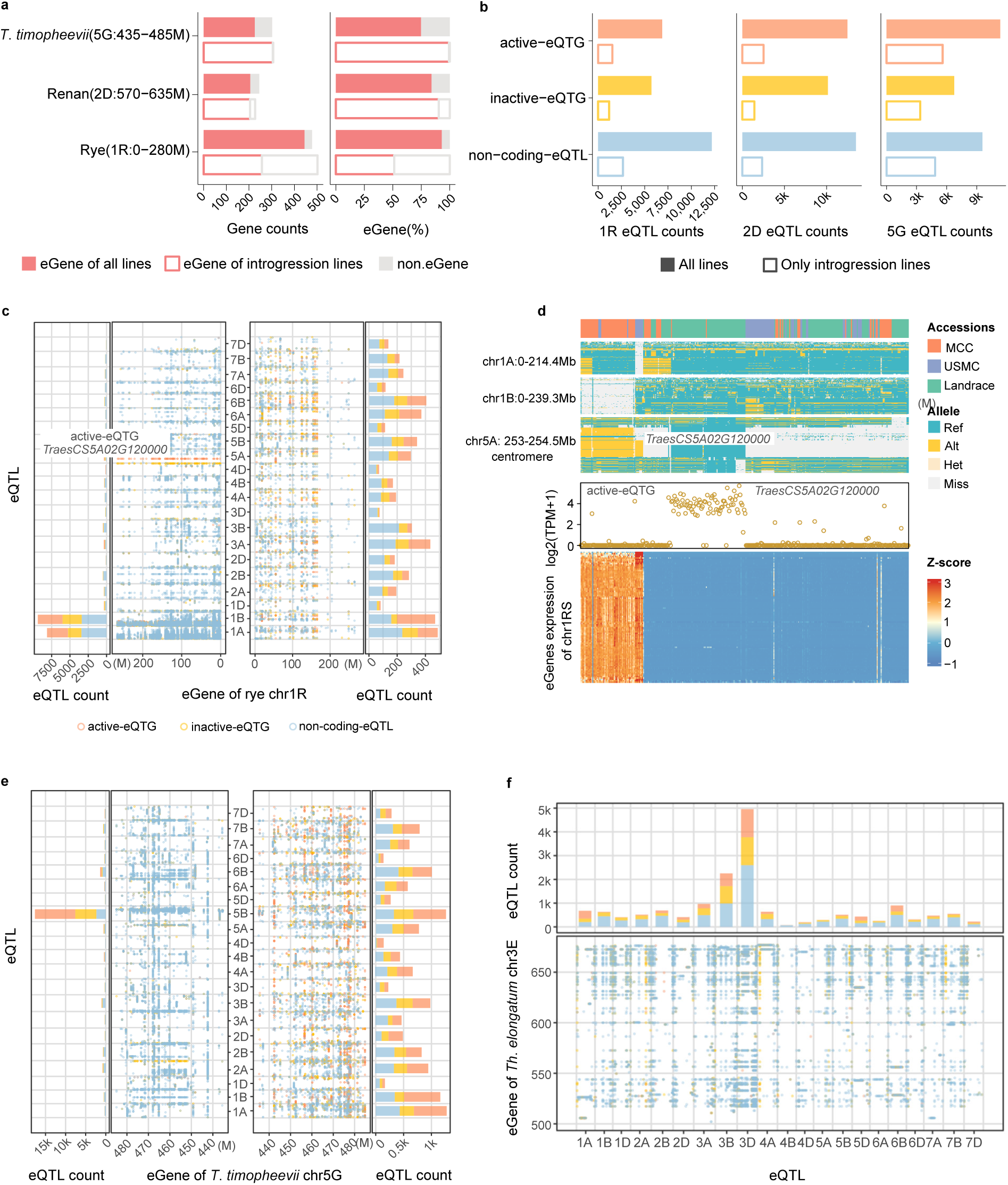
The eQTLs map for distant introgression calculated using all wheat lines of the RNA-seq panel. **a** The number of eGenes calculated for all lines within the introgression regions of rye (1R: 0-280M), Renan (2D: 570-635M), and *T*. *timopheevii* (5G: 435-485M). **b** The number of eQTLs corresponding to the eGenes calculated for all lines of the panel. **c** The eQTL map of introgressed genes on rye chr1RS. The Y-axis represents the position of SNP on wheat chromosomes, the dot plot X-axis represents the position of introgressed eGenes on chr1RS, and the X-axis of the bar plot represents the number of eQTLs. **d** Hotspot of active-eQTG on chr5A in panel c: *TraesCS5A02G120000*. The color bar at the top represents subpopulation. The three heatmaps show the regions of 1AS, 1BS, and the 5A interval containing *TraesCS5A02G120000*. The hollow scatter plot shows the expression of *TraesCS5A02G120000* across all lines, and the heatmap at the bottom shows the expression profiles of the 1RS eGenes potentially regulated by *TraesCS5A02G120000*. **e** The eQTL map of chr5G *T*. *timopheevii* introgression. The other annotations are the same as in panel c. **f** The eQTL map of *Th*. *elongatum* chr3E introgression.

**Supplementary Fig. 9.**
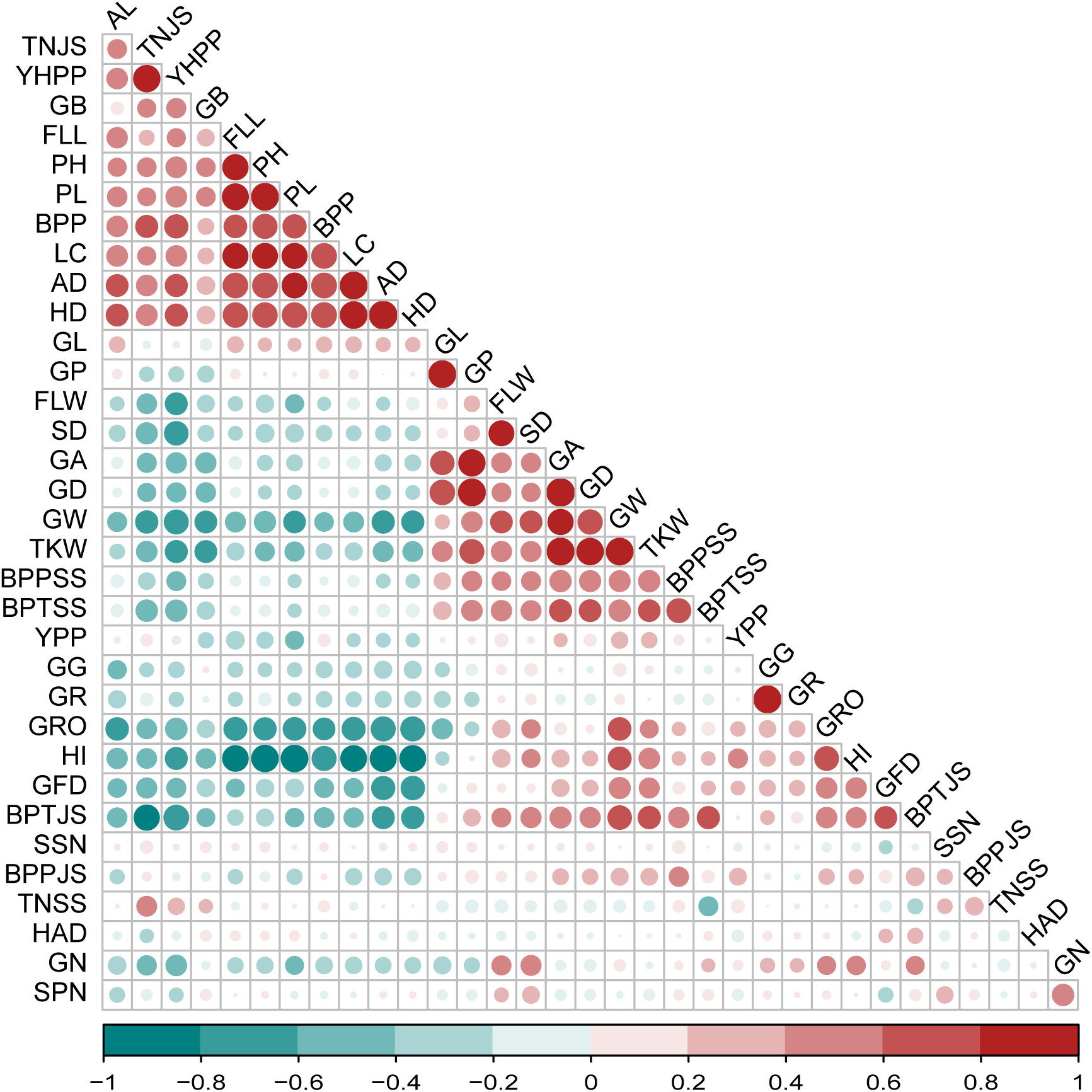
Pearson correlation coefficient plot of 34 agronomic trait phenotypes.

**Supplementary Fig. 10.**
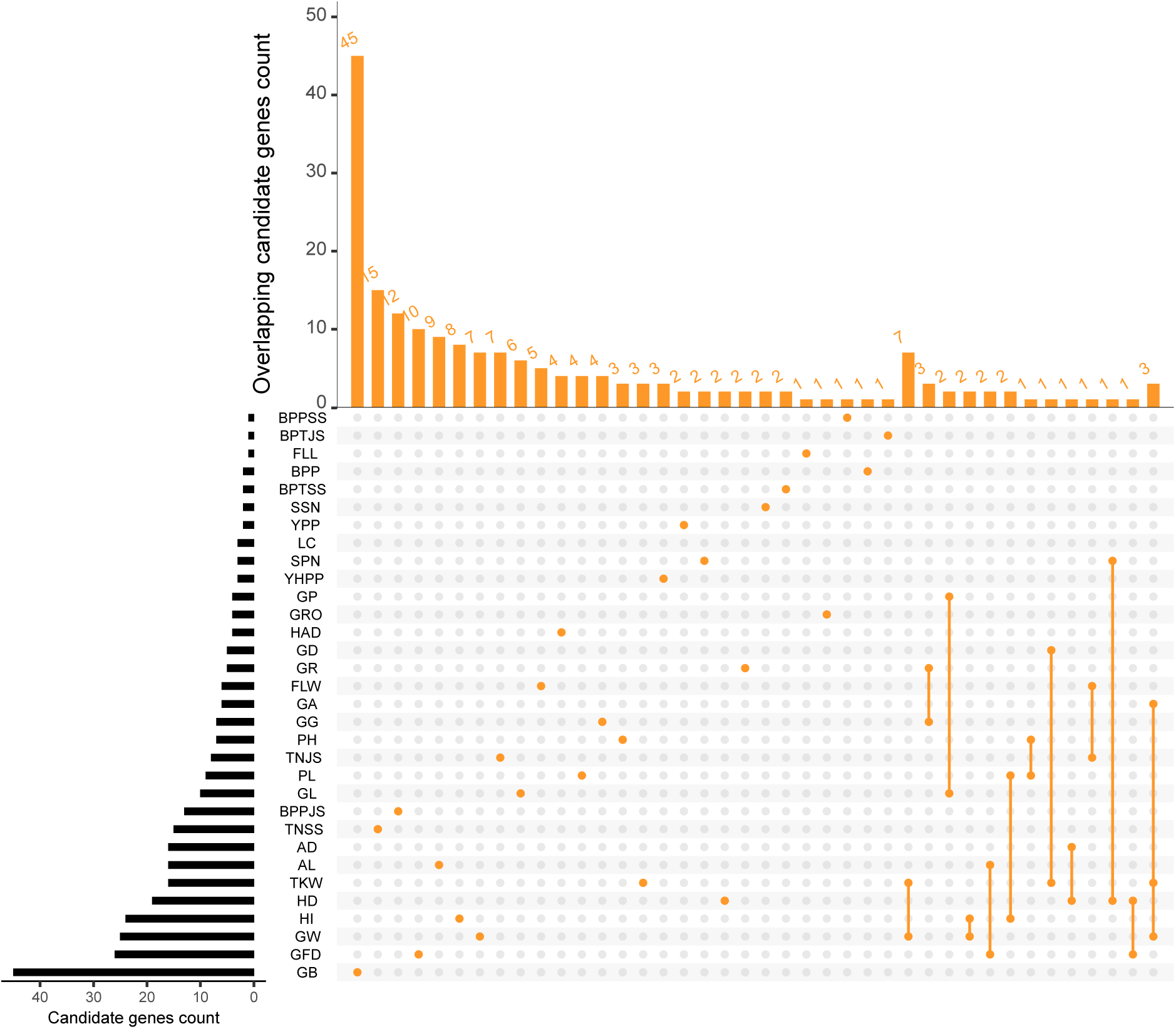
Upset plot of candidate genes for 34 field agronomic traits. Traits without candidate genes are not shown in the plot.

**Supplementary Fig. 11.**
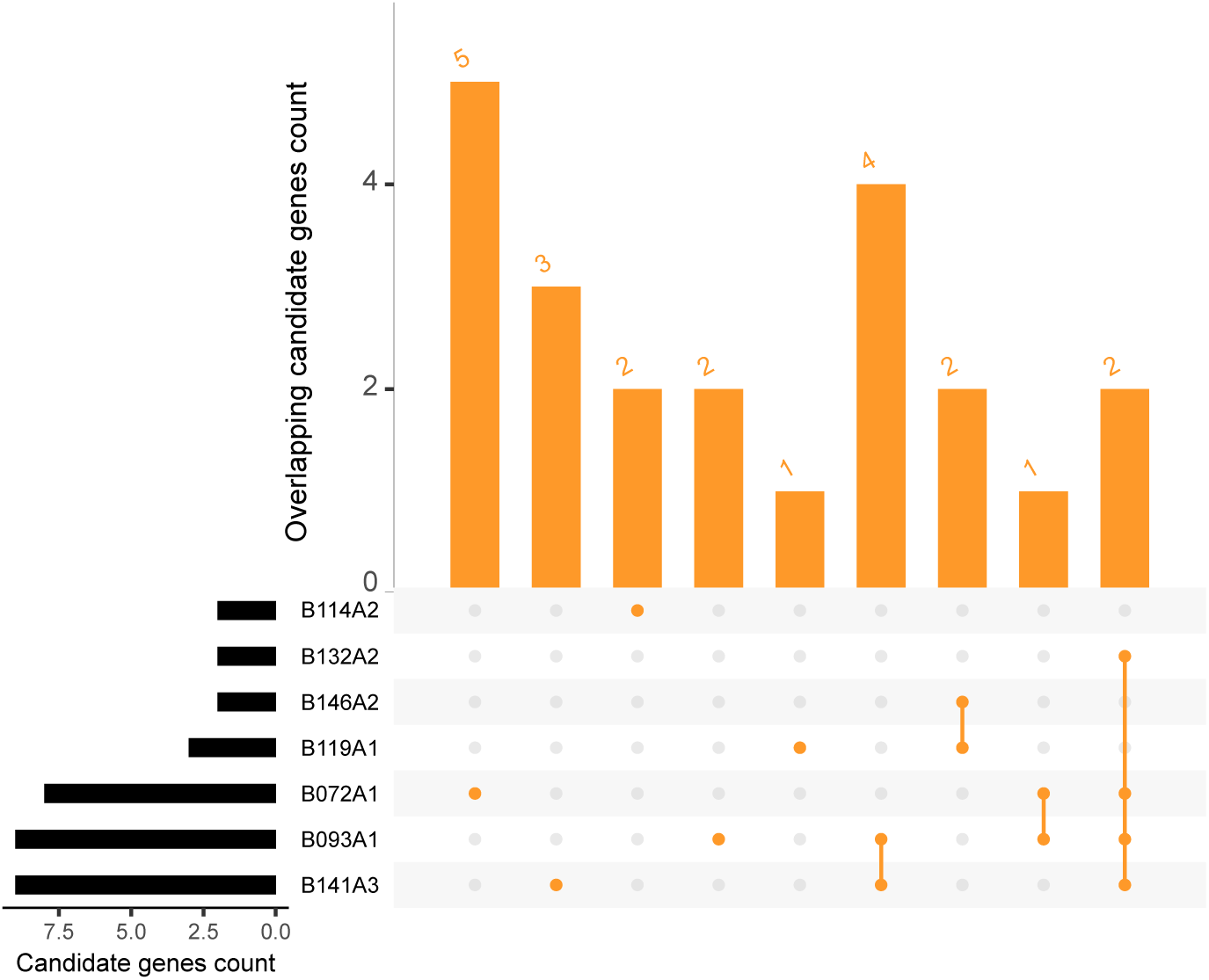
Upset plot of candidate genes for seedling-stage resistance to 8 powdery mildew isolates. Traits without candidate genes are not shown in the plot.

**Supplementary Fig. 12.**
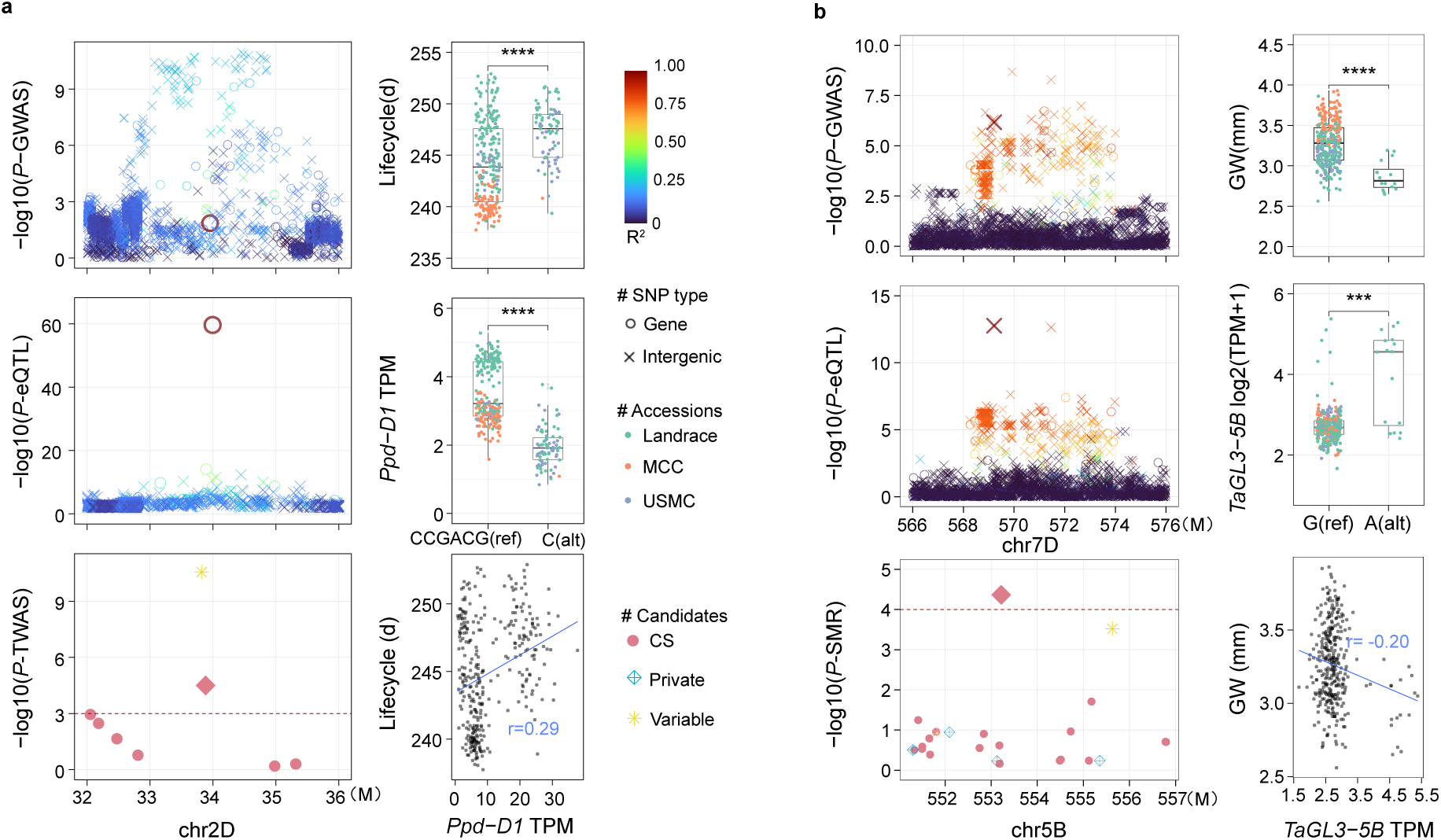
Identification of candidate genes using a combination of different methods. **a** The cloned photoperiod gene *Ppd*-*D1* identified by combining GWAS, eQTL and TWAS. **b** The cloned grain size gene *TaGL3*-*5B* identified by combining GWAS, eQTL, and SMR.

**Supplementary Fig. 13.**
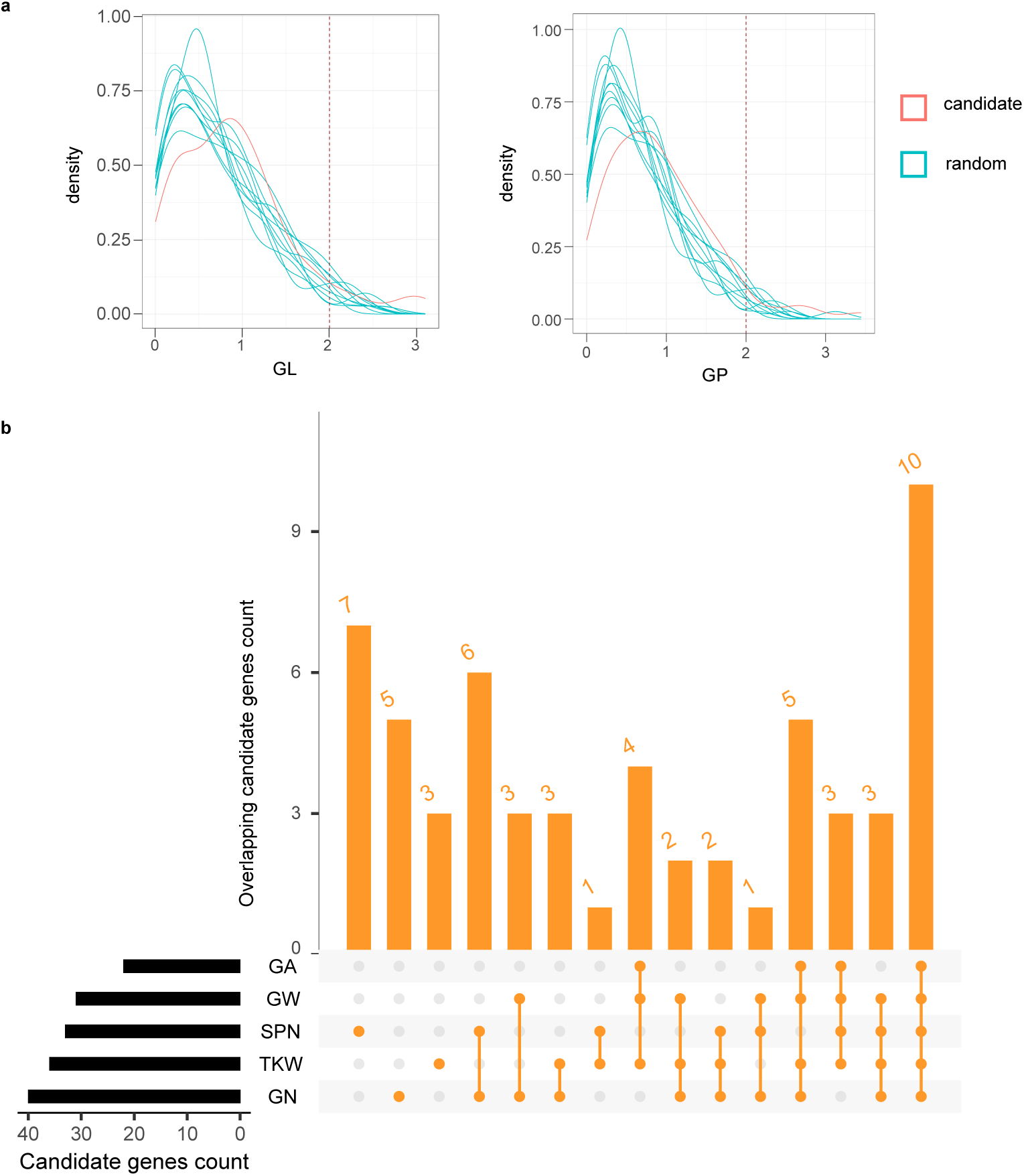
Significant candidate genes for 5 agronomic traits validated using the KN9204 mutant library. **a** The phenotypes of mutants and wild-type for the predicted candidate genes were analyzed using a two-tailed Student’s *t*-test, with the red line representing the *t*-test distribution for the candidate genes. Randomly selecting genes 10 times, with each selection containing the same number of genes as the candidate genes, was verified using the KN9204 mutant library, with the blue-green line representing the *t*-test distribution for the randomly selected genes. **b** Significant candidate genes for 5 agronomic traits validated using the KN9204 mutant library.

**Supplementary Fig. 14.**
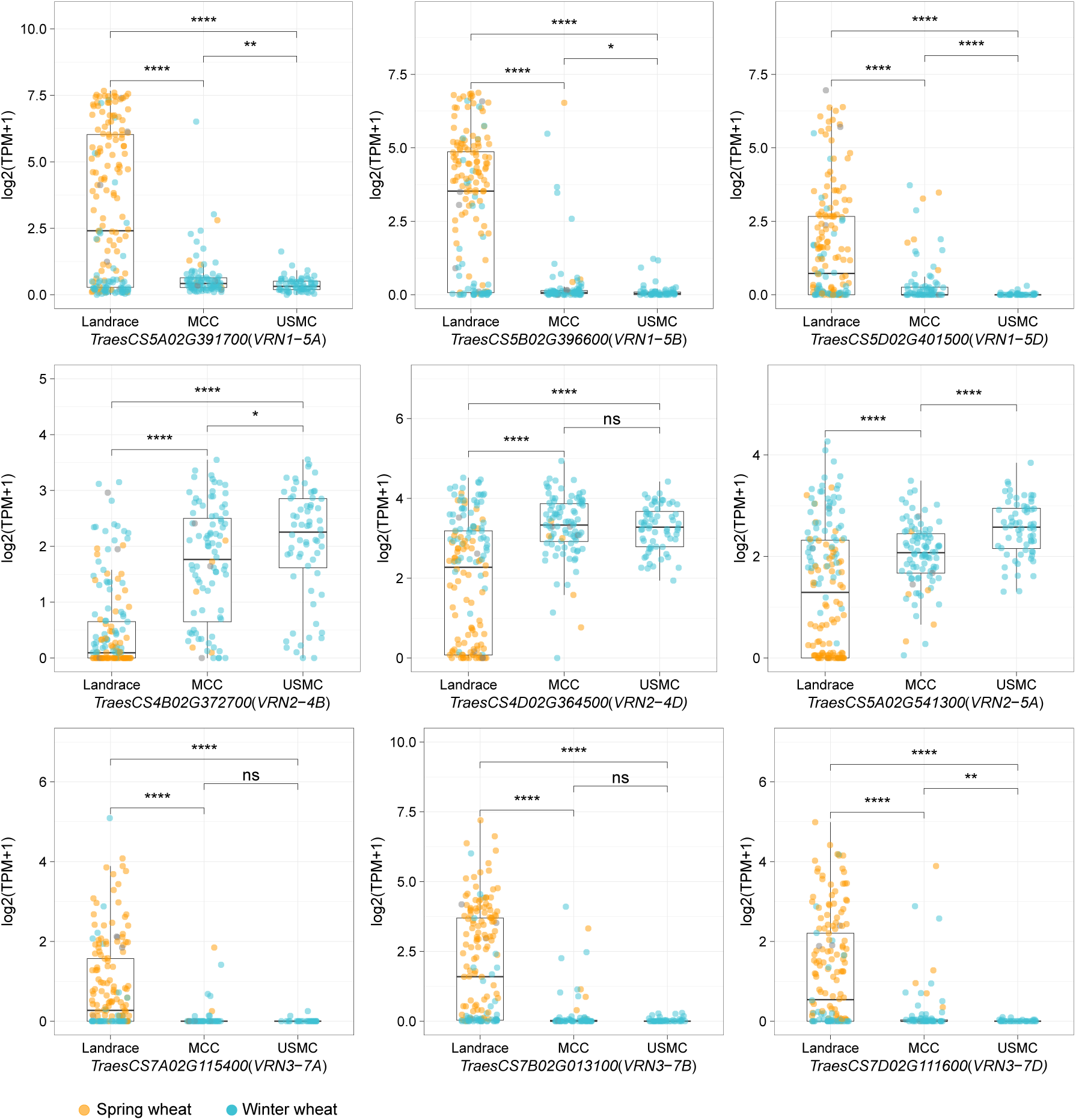
Expression of *VRN1*, *VRN2*, and *VRN3* in landraces, MCC(Modern Chinese Cultivar) and USMC(United States Modern Cultivar). Orange points represent spring wheat, and blue-green points represent winter wheat.

**Supplementary Fig. 15.**
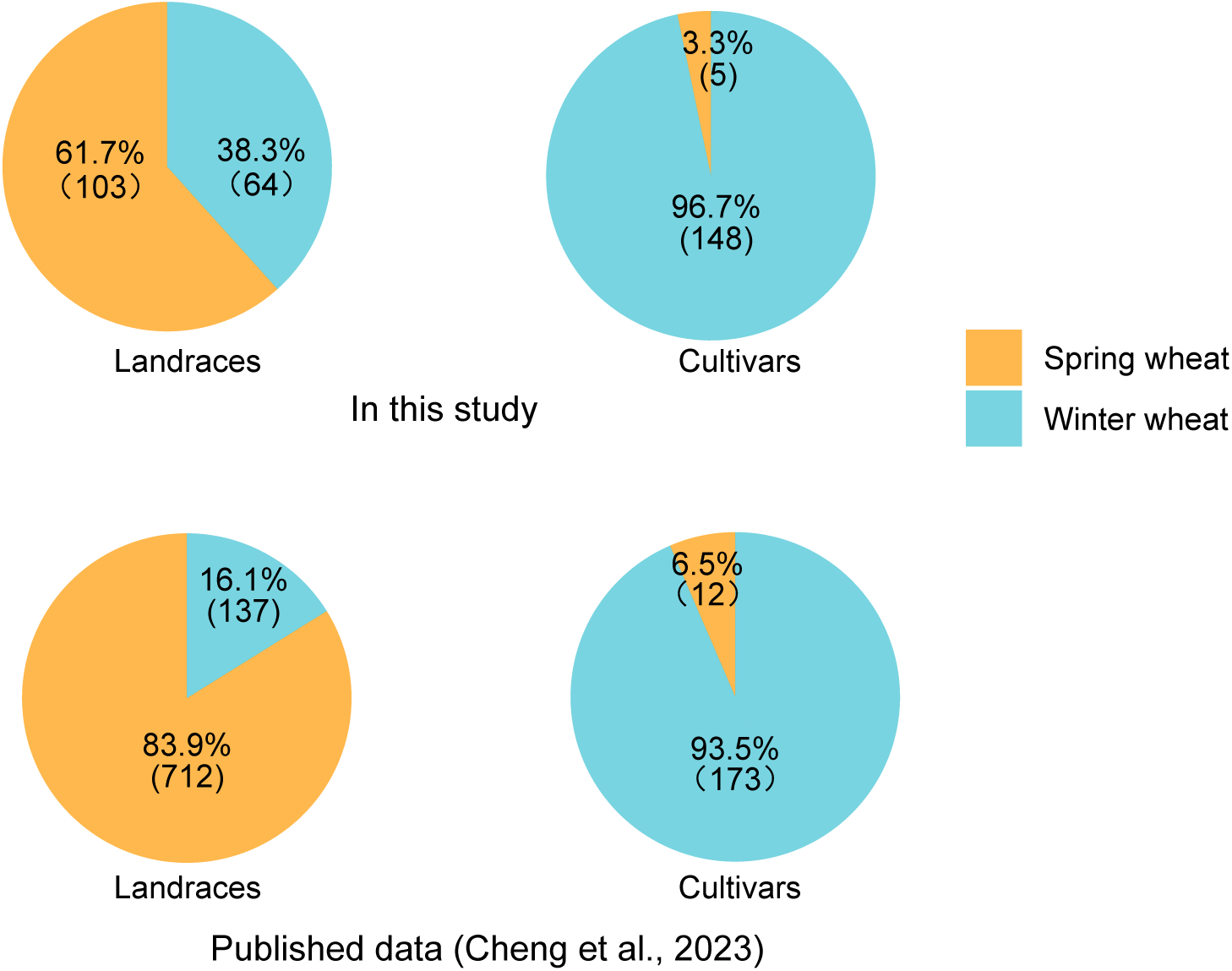
The proportion of spring wheat and winter wheat in landraces and cultivars in this study and previously published data.

**Supplementary Fig. 16.**
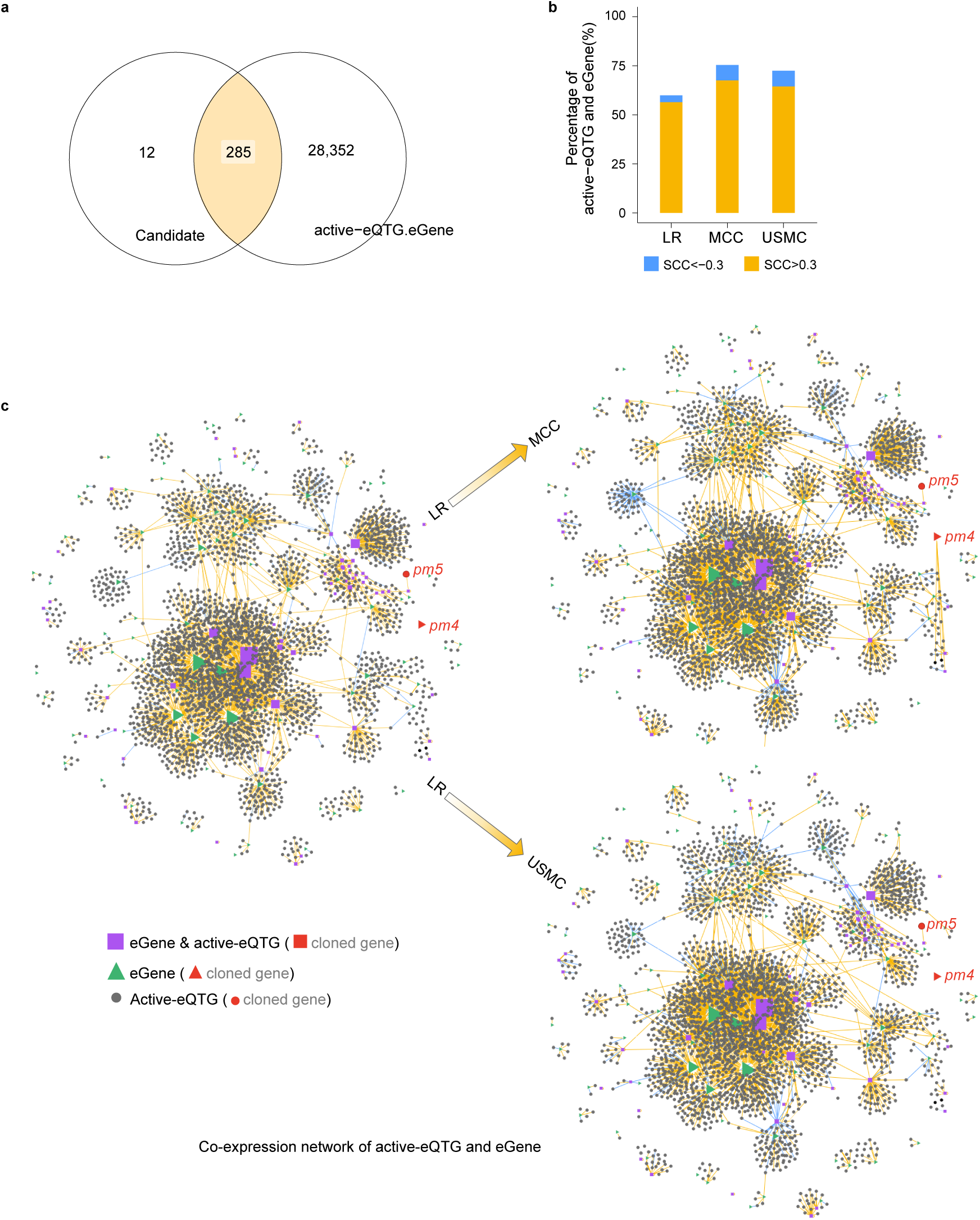
Co-expression regulatory network of candidate genes impacted by breeding selection. **a** Venn diagram of powdery mildew resistance candidate genes (predicted by one method), and eGenes regulated by active-eQTGs. **b** The proportion of candidate genes in the orange blocks of panel a and active-eQTGs with Spearman’s rank Correlation Coefficient |SCC| > 0.3 in different sub-populations. **c** Co-expression regulatory network of the genes in the orange blocks from panel a and active-eQTGs in different sub-populations. The orange lines represent the gene expression correlation (SCC>0.3) between eGenes and active-eQTGs in the sub-populations, and the blue lines represent the negative correlation (SCC< −0.3) between eGenes and active-eQTGs. eGenes and active-eQTGs with no link represent gene expression correlations where |SCC| < 0.3 in the sub-populations.

## Supplementary Tables

**Supplementary Table 1** Information for wheat accessions used in the study.

**Supplementary Table 2** Distant exogenous introgression segments and alternative reference segments.

**Supplementary Table 3** RNA-seq mapping rate with different reference gene sets (%).

**Supplementary Table 4** Number of genes expressed in the Pan-gene atlas (TPM>0.5).

**Supplementary Table 5** The proportion of expressed genes of different sources in the pan-gene atlas (TPM>0.5).

**Supplementary Table 6** RNAseq of *T*. *ponticum*, Rye, Renan, *T*. *timopheevi* used in the study.

**Supplementary Table 7** Primers of gDNA and qRT-PCR for verifying the status of introgression expression.

**Supplementary Table 8** The results of qRT-PCR in rye and the 1RS.1BL translocation line.

**Supplementary Table 9** The information of 8 *Bgt* isolates used for phenotyping the wheat panel.

**Supplementary Table 10** The QTLs of 34 field agronomic traits based on GWAS.

**Supplementary Table 11** The candidate genes of 34 agronomic traits based on TWAS/FUSION (Functional Summary-based Imputation).

**Supplementary Table 12** The candidate genes of 34 agronomic traits based on SMR (Summary-data-based Mendelian Randomization).

**Supplementary Table 13** The candidate genes of 34 agronomic traits using SCC (Spearman Correlation Coefficients).

**Supplementary Table 14** High-confidence candidate genes for 34 agronomic traits identified by at least two methods.

**Supplementary Table 15** The QTLs of 8 *Bgt* isolates based on GWAS.

**Supplementary Table 16** The candidate genes of 8 *Bgt* isolates based on TWAS/FUSION (Functional Summary-based Imputation).

**Supplementary Table 17** The candidate genes of 8 *Bgt* isolates based on SMR (Summary-data-based Mendelian Randomization).

**Supplementary Table 18** The candidate genes of 8 *Bgt* isolates based on SCC (Spearman Correlation Coefficients).

**Supplementary Table 19** High-confidence candidate genes for 8 *Bgt* isolates identified by at least two methods.

**Supplementary Table 20** Validation of candidate genes using the indexed KN9204 mutant library by comparing trait difference between wild type lines and at least 5 mutant lines.

**Supplementary Table 21** Mutation information of validated candidate genes in the indexed KN9204 mutant library.

**Supplementary Table 22** Mutant phenotype data of validated candidate genes in the indexed KN9204 mutant library.

**Supplementary Table 23** Spearman correlation coefficients of eGene and active-eQTG expression in LR and MCC for candidate genes of 34 agronomic traits.

**Supplementary Table 24** Spearman Correlation Coefficients of eGene and active-eQTG expression in LR and USMC for candidate genes of 34 field agronomic traits.

**Supplementary Table 25** Spearman Correlation Coefficients of eGene and active-eQTG expression in LR, MCC and USMC for candidate genes of powdery mildew resistance.

