## Supplementary Note for "Population-scale gene expression analysis reveals the contribution of expression diversity to the modern wheat improvement"

**Supplemental Note 1**

We integrated the high-confidence genes from Rye, Renan, and *Thinopyrum*. *elongatum* (with *Th. elongatum* as a substitute for *T. ponticum*, which lacks a published genome) and Chinese Spring into a comprehensive reference gene atlas. We observed that wheat lines with deletions at chr1A:0-214.4M and chr1B:0-239.3M have fewer quantified genes on chromosomes 1A and 1B, but show a higher number of quantified genes in the Rye: (chr1R:0-280M) region. Consequently, the Rye: (chr1RS:0-280M) region provides a more accurate quantification of gene expression in these deletion lines. Similarly, wheat lines with deletions at CS: (chr2D: 570.1-619.2M) exhibit fewer quantified genes on CS: chr2D compared to non-deletion lines, but show more quantified genes in Renan: (chr2D:575-630M). This suggests that these lines may harbor the same introgression in this region as Renan2D, potentially originating from *A. markgrafii* [27]. Therefore, Renan: (chr2D:575-630M) offers more accurate gene quantification for these lines. Wheat lines with deletions at chr3D:500M-615,552,423 display fewer quantified genes on CS: chr3D but show the opposite trend in *Th. elongatum*: (chr3E:500M-676,969,686), indicating that this deletion corresponds to a known *T. ponticum* introgression. Thus, *Th. elongatum*: (chr3E:500M-676,969,686) provides a more precise estimate of gene expression for these lines. Similarly, wheat lines with deletions at chr5B:497.2-537M have fewer quantified genes on chr5B but exhibit higher gene expression in *T. timopheevi*: (chr5G: 440-485M), suggesting that this region may represent a *T. timopheevi* introgression. Therefore, *T. timopheevi*: (chr5G: 440-485M) offers a more accurate quantification of gene expression in these lines (Fig. 1d). We observed that using genomes with potential introgressions led to more accurate gene expression quantification for deletion lines, although gene expression was also observed outside the introgressed regions. To minimize these effects, we focused on specific introgressed regions for further analysis: Rye: (chr1RS: 0-280M), Renan: (chr2D: 575-630M), *Th. elongatum*: (chr3E: 500M-676,969,686) and *T. timopheevi*: (chr5G: 440-485M).

**Supplemental Note 2**

Among the eight wheat lines with the 1RS.1AL introgression, there were 543 genes (43.5%) that had TPM > 0.5 in at least one sample. For the 52 wheat lines with the 1RS.1BL introgression, 518 genes (41.5%) satisfied the criterion of TPM > 0.5 in three samples. In the 50 Renan2D reference introgression lines, 254 genes (31.7%) met the criterion of having TPM > 0.5 in three samples. The four lines with *Th. elongatum* introgression showed that 510 genes (24.0%) had TPM > 0.5 in one sample. Lastly, among the 75 introgression lines with *T. timopheevi* on chromosome 5B, 330 genes (23.2%) met the criterion of having TPM > 0.5 in three samples (Supplementary Fig. 1).

**Supplemental Note 3**

We found that candidate genes for five agronomic traits----Spikelet Number (SPN), Grain Number (GN), Grain Area (GA), Grain Width (GW), and Thousand Kernel Weight (TKW)----exhibited greater phenotypic variation compared to randomly selected control genes. In contrast, for the other two agronomic traits----Grain Perimeter (GP) and Grain Length (GL)----the phenotypic variation distributions of candidate genes and random control genes were almost identical. Based on the differences in distribution between candidate and random genes, significance thresholds were determined. For TKW, GA, and GW, a two-tailed Student's *t*-test *p*-value < 0.01 was considered indicative of significant phenotypic variation caused by candidate gene mutations. For SPN and GN, the two-tailed Student's *t*-test *p*-value < 0.05 was used to denote significant phenotypic variation.
